# OGT binds a conserved C-terminal domain of TET1 to regulate TET1 activity and function in development

**DOI:** 10.1101/253872

**Authors:** Joel Hrit, Cheng Li, Elizabeth Allene Martin, Eric Simental, Mary Goll, Barbara Panning

**Affiliations:** Department of Biochemistry and Biophysics, University of California San Francisco, San Francisco, CA, United States; TETRAD Graduate Program, University of California San Francisco, San Francisco, CA, United States; Developmental Biology Program, Memorial Sloan Kettering Cancer Center, New York, NY, United States; Program in Biochemistry and Structural Biology, Cell and Developmental Biology, and Molecular Biology, Weill Cornell Graduate School of Medical Sciences, Cornell University, New York, NY, United States

## Abstract

TET enzymes convert 5-methylcytosine to 5-hydroxymethylcytosine and higher oxidized derivatives. TETs stably associate with and are post-translationally modified by the nutrient-sensing enzyme OGT, suggesting a connection between metabolism and the epigenome. Here, we show for the first time that modification by OGT enhances TET1 activity *in vitro.* We identify a domain of TET1 responsible for binding to OGT and report a point mutation that disrupts the TET1-OGT interaction. We show that the TET1-OGT interaction is necessary for TET1 to rescue hematopoetic stem cell production in tet mutant zebrafish embryos, suggesting that OGT promotes TET1’s function during development. Finally, we show that disrupting the TET1-OGT interaction in mouse embryonic stem cells changes the abundance of TET-containing high molecular weight complexes and causes widespread gene expression changes. These results link metabolism and epigenetic control, which may be relevant to the developmental and disease processes regulated by these two enzymes.

## Introduction

Methylation at the 5’ position of cytosine in DNA is a widespread epigenetic regulator of gene expression. Proper deposition and removal of this mark is indispensable for normal vertebrate development, and misregulation of DNA methylation is a common feature in many diseases [1,2]. The discovery of the Ten-Eleven Translocation (TET) family of enzymes, which iteratively oxidize 5-methylcytosine (5mC) to 5-hydroxymethylcytosine (5hmC), 5-formylcytosine (5fC), and 5-carboxylcytosine (5caC), has expanded the epigenome [3-7]. These modified cytosines have multiple roles, functioning both as transient intermediates in an active DNA demethylation pathway [6,8-11] and as stable epigenetic marks [12,13] that may recruit specific readers [14].

One interesting interaction partner of TET proteins is *O*-linked N-acetylglucosamine (*O*-GlcNAc) Transferase (OGT). OGT is the sole enzyme responsible for attaching a GlcNAc sugar to serine, threonine, and cysteine residues of over 1,000 nuclear, cytoplasmic, and mitochondrial proteins [15-17]. Like phosphorylation, *O*-GlcNAcylation is a reversible modification that affects the function of target proteins. OGT’s targets regulate gene expression [18,19], metabolism [16,20,21], and signaling [22,23], consistent with OGT’s role in development and disease [24,25].

OGT stably interacts with and modifies all three TET proteins and its genome-wide distribution overlaps significantly with TETs [26-28]. Two studies in mouse embryonic stem cells (mESCs) have suggested that TET1 and OGT may be intimately linked in regulation of gene expression, as depleting either enzyme reduced the chromatin association of the other and affected expression of its target genes [26,29]. However, it is unclear to what extent these genome-wide changes are direct effects of perturbing the TET1-OGT interaction. Further work is necessary to uncover the biological importance of the partnership between TET1 and OGT.

In this work, we map the interaction between TET1 and OGT to a small C-terminal region of TET1, which is both necessary and sufficient to bind OGT. We show for the first time that OGT modifies the catalytic domain of TET1 *in vitro* and enhances its catalytic activity. We also use mutant TET1 to show that the TET1-OGT interaction promotes TET1 function in the developing zebrafish embryo. Finally,we show that in mESCs a mutation in TET1 that impairs its interaction with OGT results in alterations in gene expression, in 5mC distribution, and in the abundance of TET1-, TET2-, and TET3-containing high molecular weight complexes. Together these results suggest that OGT regulates TET1 activity, indicating that the TET1-OGT interaction may be two-fold in function - allowing TET1 to recruit OGT to specific genomic loci and allowing OGT to modulate TET1 activity.

## Materials and Methods

### Cell Culture

The mESC line LF2, and its derivatives were routinely passaged by standard methods in KO-DMEM, 10% FBS, 2 mM glutamine, 1X non-essential amino acids, 0.1 mM b-mercaptoethanol and recombinant leukemia inhibitory factor. HEK293T were cultured in DMEM, 10% FBS, and 2 mM glutamine.

### Recombinant protein purification

Full-length human OGT in the pBJG vector was transformed into BL-21 DE3 E. coli. A liquid culture was grown in LB + 50ug/mL kanamycin at 37C until OD600 reached 1.0. IPTG was added to 1mM final and the culture was induced at 16C overnight. Cells were pelleted by centrifugation and resuspended in 5mL BugBuster (Novagen) + protease inhibitors (Sigma Aldrich) per gram of cell pellet. Cells were lysed on an orbital shaker for 20 minutes at room temperature. The lysate was clarified by centrifugation at 30,000g for 30 minutes at 4C. Clarified lysate was bound to Ni-NTA resin (Qiagen) at 4C and then poured over a disposable column. The column was washed with 6 column volumes of wash buffer 1 (20mM Tris pH 8, 1mM CHAPS, 10% glycerol, 5mM BME, 10mM imidazole, 250mM NaCl) followed by 6 column volumes of wash buffer 2 (wash buffer 1 with 50mM imidazole). The protein was eluted in 4 column volumes of elution buffer (20mM Tris pH 8, 1mM CHAPS, 5mM BME, 250mM imidazole, 250mM NaCl). Positive fractions were pooled and dialyzed into storage buffer (20mM Tris pH 8, 1mM CHAPS, 0.5mM THP, 10% glycerol, 150mM NaCl, 1mM EDTA), flash frozen in liquid nitrogen and stored at -80C in small aliquots.

Mouse TET1 catalytic domain (aa1367-2039) was expressed in sf9 insect cells according to the Bac-to-Bac Baculovirus Expression System. Constructs were cloned into the pFastBac HTA vector and transformed in DH10Bac E. coli for recombination into a bacmid. Bacmid containing the insert was isolated and used to transfect adherent sf9 cells for 6 days at 25C. Cell media (P1 virus) was isolated and used to infect 20mL of sf9 cells in suspension for 3 days. Cell media (P2 virus) was isolated and used to infect a larger sf9 suspension culture for 3 days. Cells were pelleted by centrifugation, resuspended in lysis buffer (20mM Tris pH 8, 1% Triton, 10% glycerol, 20mM imidazole, 50mM NaCl, 1mM MgCl2, 0.5mM TCEP, protease inhibitors, 2.5U/mL benzonase), and lysed by douncing and agitation at 4C for 1 hour. The lysate was clarified by centrifugation at 48,000g for 30 minutes at 4C and bound to Ni-NTA resin (Qiagen) at 4C, then poured over a disposable column. The column was washed with 5 column volumes of wash buffer (20mM Tris pH 8, 0.3% Triton, 10% glycerol, 20mM imidazole, 250mM NaCl, 0.5mM TCEP, protease inhibitors). The protein was eluted in 5 column volumes of elution buffer (20mM Tris pH 8, 250mM imidazole, 250mM NaCl, 0.5mM TCEP, protease inhibitors). Positive fractions were pooled and dialyzed overnight into storage buffer (20mM Tris pH 8, 150mM NaCl, 0.5mM TCEP). Dialyzed protein was purified by size exclusion chromatography on a 120mL Superdex 200 column (GE Healthcare). Positive fractions were pooled, concentrated, flash frozen in liquid nitrogen and stored at -80C in small aliquots.

### Overexpression in HEK293T cells and immunoprecipitation

Mouse Tet1 catalytic domain (aa1367-2039) and truncations and mutations thereof were cloned into the pcDNA3b vector. GFP fusion constructs were cloned into the pcDNA3.1 vector. Human OGT constructs were cloned into the pcDNA4 vector. Plasmids were transiently transfected into adherent HEK293T cells at 70-90% confluency using the Lipofectamine 2000 transfection reagent (ThermoFisher) for 1-3 days.

Full-length mouse Tet1 and mutations thereof were cloned into the pCAG vector. Plasmids were transiently transfected into adherent HEK293T cells at 70-90% confluency using the PolyJet transfection reagent (SignaGen) for 1-3 days.

Transiently transfected HEK293T cells were harvested, pelleted, and lysed in IP lysis buffer (50mM Tris pH 8, 200mM NaCl, 1% NP40, 1x HALT protease/phosphatase inhibitors). For pulldown of FLAG-tagged constructs, cell lysate was bound to anti-FLAG M2 magnetic beads (Sigma Aldrich) at 4C. For pulldown of GFP constructs, cell lysate was bound to magnetic protein G dynabeads (ThermoFisher) conjugated to the JL8 GFP monoclonal antibody (Clontech) at 4C. Beads were washed 3 times with IP wash buffer (50mM Tris pH 8, 200mM NaCl, 0.2% NP40, 1x HALT protease/phosphatase inhibitors). Bound proteins were eluted by boiling in SDS sample buffer.

### *In vitro* transcription/translation and immunoprecipitation

GFP fused to TET C-terminus peptides were cloned into the pcDNA3.1 vector and transcribed and translated *in vitro* using the TNT Quick Coupled Transcription/Translation System (Promega).

For immunoprecipitation, recombinant His-tagged OGT was coupled to His-Tag isolation dynabeads (ThermoFisher). Beads were bound to *in vitro* translation extract diluted 1:1 in binding buffer (40mM Tris pH 8, 200mM NaCl, 40mM imidazole, 0.1% NP40) at 4C. Beads were washed 3 times with wash buffer (20mM Tris pH 8, 150mM NaCl, 20mM imidazole, 0.1% NP40). Bound proteins were eluted by boiling in SDS sample buffer.

### Recombinant protein binding assay

20uL reactions containing 2.5uM rOGT and 2.5uM rTET1 CD wt or D2018A were assembled in binding buffer (50mM Tris pH 7.5, 100mM NaCl, 0.02% Tween-20) and pre-incubated at room temperature for 15 minutes. TET1 antibody (Millipore 09-872) was bound to magnetic Protein G Dynabeads (Invitrogen), and beads added to reactions following pre-incubation. Reactions were bound to beads for 10 minutes at room temperature. Beads were washed 3 times with 100uL binding buffer, and bound proteins were recovered by boiling in SDS sample buffer and analyzed by SDS-PAGE and coomassie stain.

### Western blots

For western blot, proteins were separated on a denaturing SDS-PAGE gel and transferred to PVDF membrane. Membranes were blocked in PBST + 5% nonfat dry milk at room temp for >10 minutes or at 4C overnight. Primary antibodies used for western blot were: FLAG M2 monoclonal antibody (Sigma Aldrich F1804), OGT polyclonal antibody (Santa Cruz sc32921), OGT monoclonal antibody (Cell Signaling D1D8Q), His6 monoclonal antibody (Thermo MA1-21315), JL8 GFP monoclonal antibody (Clontech), and *O*-GlcNAc RL2 monoclonal antibody (Abcam). Secondary antibodies used were goat anti-mouse HRP and goat anti-rabbit HRP from BioRad. Blots were incubated with Pico Chemiluminescent Substrate (ThermoFisher) and exposed to film in a dark room.

### Slot blot

Prior to dilution of genomic DNA samples, biotinylated *E. coli* gDNA was added as a loading control (see below). DNA samples were denatured in 400mM NaOH + 10mM EDTA by heating to 95C for 10 minutes. Samples were placed on ice and neutralized by addition of 1 volume of cold NH4OAc pH 7.2. DNA was loaded onto a Hybond N+ nylon membrane (GE) by vacuum using a slot blot apparatus. The membrane was dried at 37C and DNA was covalently linked to the membrane by UV crosslinking (700uJ/cm^2^ for 3 minutes). Antibody binding and signal detection were performed as outlined for western blotting. Primary antibodies used were 5mC monoclonal antibody (Active Motif 39649) and 5hmC monoclonal antibody (Active Motif 39791).

For the loading control, membranes were analyzed using the Biotin Chromogenic Detection Kit (Thermo Scientific) according to the protocol. Briefly, membranes were blocked, probed with streptavidin conjugated to alkaline phosphatase (AP), and incubated in the AP substrate BCIP-T (5-bromo-4-chloro-3-indolyl phosphate, p-toluidine salt). Cleavage of BCIP-T causes formation of a blue precipitate.

For quantification of slot blots, at least 3 biological replicates were used. Signal was normalized to the loading control and significance was determined using the unpaired t test.

### Preparation of lambda DNA substrate

Linear genomic DNA from phage lambda (dam-, dcm-) containing 12bp 5’ overhangs was purchased from Thermo Scientific. Biotinylation was performed by annealing and ligating a complementary biotinylated DNA oligo. Reactions containing 175ng/uL lambda DNA, 2uM biotinylated oligo, and 10mM ATP were assembled in 1x T4 DNA ligase buffer, heated to 65C, and cooled slowly to room temperature to anneal. 10uL T4 DNA ligase was added and ligation was performed overnight at room temperature. Biotinylated lambda DNA was purified by PEG precipitation. To a 500uL ligation reaction, 250uL of PEG8000 + 10mM MgCl_2_ was added and reaction was incubated at 4C overnight with rotation. The next day DNA was pelleted by centrifugation at 14,000g at 4C for 5 minutes. Pellet was washed with 1mL of 75% ethanol and resuspended in TE.

Biotinylated lambda DNA was methylated using M.SssI CpG methyltransferase from NEB. 20uL reactions containing 500ng lambda DNA, 640uM S-adenosylmethionine, and 4 units methyltransferase were assembled in 1x NEBuffer 2 supplemented with 20mM Tris pH 8 and incubated at 37C for 1 hour. Complete methylation was confirmed by digestion with the methylation-sensitive restriction enzyme BstUI from NEB.

### *In vitro* TET1 CD *O*-GlcNAcylation

*In vitro* modification of rTET1 CD with rOGT was performed as follows: 10uL reactions containing 1uM rTET1 CD, 1-5uM rOGT, and 1mM UDP-GlcNAc were assembled in reaction buffer (50mM HEPES pH 6.8, 150mM NaCl, 10% glycerol, 0.5mM TCEP) and incubated at 37C for 30-60 minutes or at 4C for 18-24 hours.

### *In vitro* TET1 CD activity assays

20uL reactions containing 100ng biotinylated, methylated lambda DNA, rTET1 CD (from frozen aliquots or from *in vitro O*-GlcNAcylation reactions), and TET cofactors (1mM alpha-ketoglutarate, 2mM ascorbic acid, 100uM ferrous ammonium sulfate) were assembled in reaction buffer (50mM HEPES pH 6.8, 100mM NaCl) and incubated at 37C for 10-60 minutes. Reactions were stopped by addition of 1 volume of 2M NaOH + 50mM EDTA and DNA was analyzed by slot blot.

### Generation of mouse embryonic stem cell lines

mESC lines (Fig. 3 – figure supplement 1A, B) were derived using CRISPR-Cas9 genome editing. A guide RNA to the Tet1 3’UTR was cloned into the px459-Cas9-2A-Puro plasmid using published protocols [30] with minor modifications. Templates for homology directed repair were amplified from Gene Blocks (IDT) (Supplementary File 1A, B). Plasmid and template were co-transfected into LF2 mESCs using FuGENE HD (Promega) according to manufacturer protocol. After two days cells were selected with puromycin for 48 hours, then allowed to grow in antibiotic-free media. Cells were monitored for green or red fluorescence (indicating homology directed repair) and fluorescent cells were isolated by FACS 1-2 weeks after transfection. All cell lines were propagated from single cells and correct insertion was confirmed by PCR genotyping (Fig. 3 – figure supplement 1A, B, Supplementary File 1A).

### Chromosome Spread Preparations

Chromosome spreads were prepared using hypotonic swelling and methanol:acetic acid fixation following established protocols[31]

### Immunofluorescence and Imaging

Slides were incubated in 1M HCl at 37C for 45 minutes to denature chromatin, then neutralized in 100mM Tris pH 7.6 at room temperature for 10 minutes. Slides were washed twice in PBST for 5 minutes, then blocked in IF blocking buffer (PBS + 5% goat serum, 0.2% fish skin gelatin, 0.2% Tween-20) at room temperature for 1 hour. Primary antibodies were diluted in blocking buffer and incubated on slides at 4C overnight. Primary antibodies used were FLAG M2 monoclonal antibody (Sigma Aldrich F1804), 5mC monoclonal antibody (Active Motif 39649), and 5hmC polyclonal antibody (Active Motif 39791). Slides were washed twice in PBST for 5 minutes, then incubated with secondary antibodies diluted in IF blocking buffer. Secondary antibodies used were Alexa488-conjugated goat anti-rabbit IgG (Jackson 711-545-152), Cy3-conjugated goat anti-rabbit IgG (Jackson 715-165-152), and Cy3-conjugated goat anti-mouse IgG (Jackson 715-165-150). Slides were washed three times in PBST for 5 minutes with DAPI added to the second wash (final concentration 100ng/mL). Slides were then mounted using prolong gold antifade (Molecular Probes P36930) and imaged. For comparisons between cell lines, all images were taken with the same exposure time, and any image processing was performed identically on all images. 10um scale bars are included.

### mESC nuclei isolation and fractionation

For isolation of nuclei, mESCs were lysed in buffer 1 (10mM Tris pH 8, 320mM sucrose, 3mM CaCl2, 2mM MgOAc, 0.1mM EDTA, 0.1% Triton X-100, 1mM DTT, and protease inhibitors). Lysed cells were diluted in two volumes of buffer 2 (10mM Tris pH 8, 2M sucrose, 5mM MgOAc, 0.1mM EDTA, 5mM DTT, and protease inhibitors), then layered over buffer 2 in a centrifuge tube. Nuclei were pelleted by centrifugation at 37,000rpm at 4C for 45 minutes and recovered.

Nuclei were lysed in nuclear lysis buffer (20mM Tris pH 8, 300mM NaCl, 10% glycerol, 0.25% Igepal, and protease inhibitors). Nuclear proteins were fractionated on a Superose 6 Increase 10/300 GL column at 0.5mL/min in nuclear lysis buffer. 0.5mL fractions were collected, concentrated, and analyzed by western blot.

### RNA-seq

Libraries for RNA-seq were prepared using the TruSeq PolyA kit. Two replicates of Tet1 wild-type and three replicates of Tet1 D2018A lines were passed quality control and were analyzed. Single-end 50bp RNAseq was performed on an Illumina HiSeq 4000 sequencer. Reads were mapped to the mouse genome (GRCm38.78), and reads uniquely mapped to known mRNAs were used to assess expression changes between genes. Only genes with FDR < 0.1 were considered in downstream analyses.

### RT-qPCR

Total RNA was isolated from mESCs using Direct-zol RNA miniprep kit from Zymo. 1ug of RNA was used for cDNA synthesis using the iScript Reverse Transcription kit from BioRad. cDNA was used for qPCR using the SensiFAST SYBR Lo-Rox kit from Bioline. Relative gene expression levels were calculated using the ΔΔC_t_ method. See Supplementary File 2A for primer sequences.

### Bisulfite conversion and analysis

Reactions containing 200ng of mESC genomic DNA + 200ng phage lambda genomic DNA were bisulfite treated and purified using the EZ DNA Methylation Lightning kit from Zymo. DMRs for the genes *H19, PeglO,* and *Mest* were amplified using bisulfite specific primers (see Supplementary File 2B for primer sequences and genomic coordinates). Amplified DMRs were cloned into the pCR-Blunt II-TOPO plasmid and sequenced. A region of phage lambda DNA was sequenced to confirm complete bisulfite conversion.

### Zebrafish mRNA rescue experiments

Zebrafish husbandry was conducted under full animal use and care guidelines with approval by the Memorial Sloan-Kettering animal care and use committee. For mRNA rescue experiments, mTET1D2018A and mTET1wt plasmids were linearized by NotI digestion. Capped RNA was synthesized using mMessage mMachine (Ambion) with T7 RNA polymerase. RNA was injected into one-cell-stage embryos derived from tet2^mk17/mk17^, tet3^mk18/+^ intercrosses at the concentration of 100pg/embryo [32]. Injected embryos were raised under standard conditions at 28.5°C until 30 hours post-fertilization (hpf) at which point they were fixed for *in situ* hybridization using an antisense probe for *runx1.* The *runx1* probe is described in [33]*; in situ* hybridization was performed using standard methods, and runx1 levels were scored across samples without knowledge of the associated experimental conditions [34]. Examples of larvae categorized as runx1 high and runx1 low are provided in Supplementary File 3. *tet2/3* double mutants were identified based on morphological criteria and mutants were confirmed by PCR genotyping after in situ hybridization using previously described primers [32].

For sample size estimation for rescue experiments, we assume a background mean of 20% positive animals in control groups. We anticipate a significant change would result in at least a 30% difference between the experimental and control means with a standard deviation of no more than 10. Using the 1-Sample Z-test method, for a specified power of 95% the minimum sample size is 4. Typically, zebrafish crosses generate far more embryos than required. Experiments are conducted using all available embryos. The experiment is discarded if numbers for any sample are below this minimum threshold when embryos are genotyped at the end of the experimental period. Injections were separately performed on clutches from five independent crosses; p values are based on these replicates and were derived from the unpaired two-tailed *t* test. Embryo numbers for all five biological replicates are included in Supplementary File 3.

For the dot blot, genomic DNA was isolated from larvae at 30hpf by phenol-chloroform extraction and ethanol precipitation. Following RNase treatment and denaturation, 2-fold serially diluted DNA was spotted onto nitrocellulose membranes. Cross-linked membranes were incubated with 0.02% methylene blue to validate uniform DNA loading. Membranes were blocked with 5% BSA and incubated with anti-5hmC antibody (1:10,000; Active Motif) followed by a horseradish peroxidase-conjugated antibody (1:15,000; Active Motif). Signal was detected using the ECL Prime Detection Kit (GE). The results of three independent experiments were quantified using ImageJ at the lowest dilution and exposure where signal was observed in Tet1 injected embryos. To normalize across blots, all values are presented as the ratio of 5hmC signal in experimental animals divided by wildtype control signal from the same blot.

### Reproducibility and Rigor

All immunostaining, IP-Westerns, and genomic DNA slot blot data are representative of at least three independent biological replicates (experiments carried out on different days with a different batch of HEK293T cells or mESCs). For targeted mESC lines, three independently derived lines for each genotype were assayed in at least two biological replicates. For *in vitro* activity and binding assays using recombinant proteins (representing multiple protein preparations), data represent at least three technical replicates (carried out on multiple days). The zebrafish rescue experiment was performed five times (biological replicates), with dot blots carried out three times. We define an outlier as a result in which all the controls gave the expected outcome but the experimental sample yielded an outcome different from other biological or technical replicates. There were no outliers or exclusions.

## Results

### A short C-terminal region of TET1 is necessary for binding to OGT

TET1 and OGT interact with each other and are mutually dependent for their localization to chromatin[26]. To understand the role of this association, it is necessary to specifically disrupt the TET1-OGT interaction. All three TETs interact with OGT via their catalytic domains [27,28,35]. We sought to identify the region within the TET1 catalytic domain (TET1 CD) responsible for binding to OGT. The TET1 CD consists of a cysteine-rich N-terminal region necessary for co-factor and substrate binding, a catalytic fold consisting of two lobes separated by a spacer of unknown function, and a short C-terminal region also of unknown function (Fig. 1A). We transiently transfected HEK293T cells with FLAG-tagged mouse TET1 CD constructs bearing deletions of each of these regions, some of which failed to express (Fig. 1B). Because HEK293T cells have low levels of endogenous OGT, we also co-expressed His-tagged human OGT (identical to mouse at 1042 of 1046 residues). TET1 constructs were immunoprecipitated (IPed) using a FLAG antibody and analyzed for interaction with OGT. We found that deletion of only the 45 residue C-terminus of TET1 (hereafter C45) prevented detectable interaction with OGT (Fig. 1B, TET1 CD del. 4). To exclude the possibility that this result is an artifact of OGT overexpression, we repeated the experiment overexpressing only TET1. TET1 CD, but not TET1 CD ΔC45, interacted with endogenous OGT, confirming that the C45 is necessary for this interaction (Fig. 1 – figure supplement 1).

**Fig. 1:**
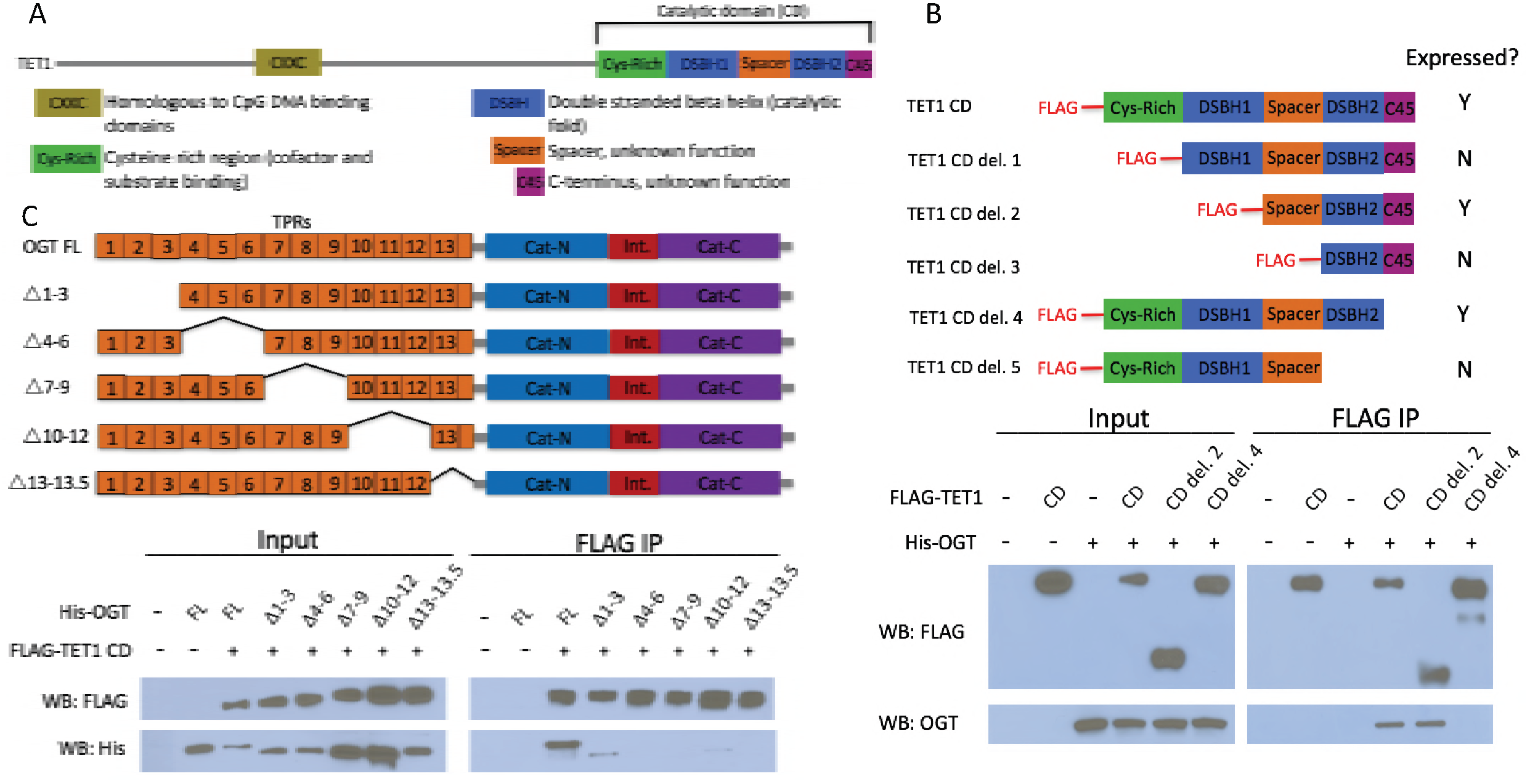
The short TET1 C1terminus is required for interaction with OGT. A) Domain architecture of TET1. B) Diagram of FLAG;tagged TET1 CD constructs expressed in HEK293T cells (upper). FLAG and OGT western blot of inputs and FLAG IPs from HEK293T cells transiently expressing FLAG;TET1 CD truncations and His;OGT (lower). C) Diagram of His;tagged OGT constructs expressed in HEK293T cells (upper). FLAG and His western blot of input and FLAG IPs from HEK293T cells transiently expressing FLAG;TET1 CD and His;OGT TPR deletions (lower).

OGT has two major domains: the N-terminus consists of 13.5 tetratricopeptide repeat (TPR) protein-protein interaction domains, and the C-terminus contains the bilobed catalytic domain (Fig. 1C). We made internal deletions of several sets of TPRs to ask which are responsible for binding to the TET1 CD. We co-transfected HEK293T cells with FLAG-TET1 CD and His6-tagged OGT constructs and performed FLAG IP and western blot as above. We found that all the TPR deletions tested impaired the interaction with TET1 CD, with deletion of TPRs 7-9, 10-12, or 13-13.5 being most severe (Fig. 1C). This result suggests that all of OGT’s TPRs may be involved in binding to the TET1 CD, or that deletion of a set of TPRs disrupts the overall structure of the repeats in a way that disfavors binding.

### Conserved residues in the TET1 C45 are necessary for the TET1-OGT interaction

An alignment of the TET1 C45 region with the C-termini of TET2 and TET3 revealed several conserved residues (Fig. 2A). We mutated clusters of three conserved residues in the TET1 C45 of FLAG-tagged TET1 CD (Fig. 2B) and co-expressed these constructs with His-OGT in HEK293T cells. FLAG pulldowns revealed that two sets of point mutations disrupted the interaction with OGT: mutation of D2018, V2021, and T2022, or mutation of V2021, T2022, and S2024 (Fig. 2C, mt1 and mt2). These results suggested that the residues between D2018 and S2024 are crucial for the interaction between TET1 and OGT. Further mutational analysis revealed that altering D2018 to A (D2018A) eliminated detectable interaction between FLAG-tagged TET1 CD and His-OGT (Fig. 2D).

**Fig. 2:**
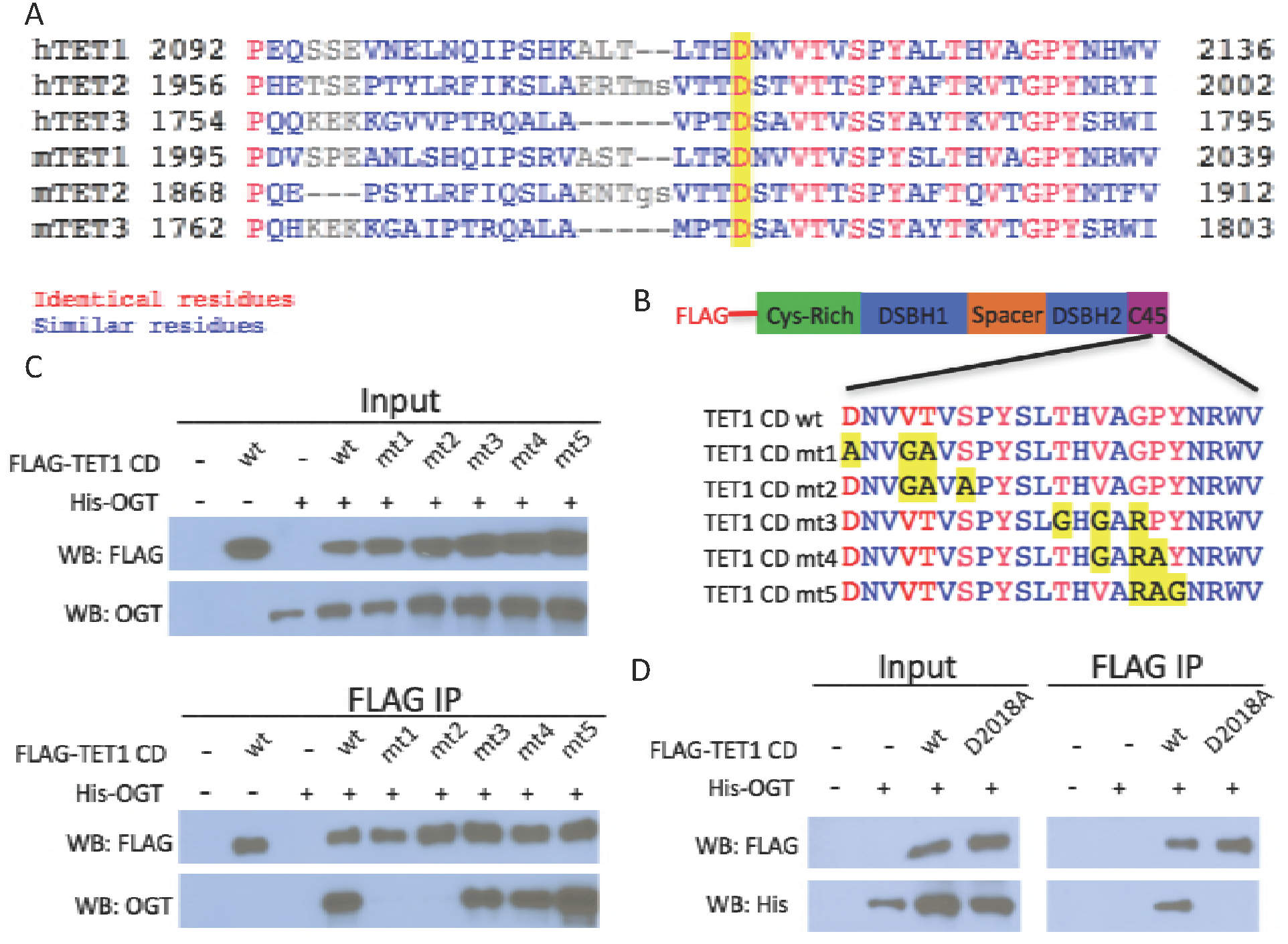
Conserved residues in the TET1 C45 are necessary for the TET1-OGT interaction. A) Alignment of the C-termini of human (h) and mouse (m) TETs 1, 2, and 3. A conserved aspartate residue mutated in D is highlighted. B) Diagram of FLAG-tagged TET1 CD constructs expressed in HEK293T cells. C) FLAG and OGT western blot of inputs and FLAG IPs from HEK293T cells transiently expressing FLAG-TET1 CD triple point mutants and His-OGT. D) FLAG and OGT western blot of inputs and FLAG IPs from HEK293T cells transiently expressing His-OGT and FLAG-TET1 CD or FLAG-TET1 CD D2018A.

### The TET1 C-terminus is sufficient for binding to OGT

Having shown that the TET1 C45 is necessary for the interaction with OGT, we next examined if it is also sufficient to bind OGT. We fused the TET1 C45 to the C-terminus of GFP (Fig. 3A) and investigated its interaction with OGT. We transiently transfected GFP or GFP-C45 into HEK293T cells and pulled down with a GFP antibody. We found that GFP-C45, but not GFP alone, bound OGT (Fig. 3B), indicating that the TET1 C45 is sufficient for interaction with OGT.

**Fig. 3:**
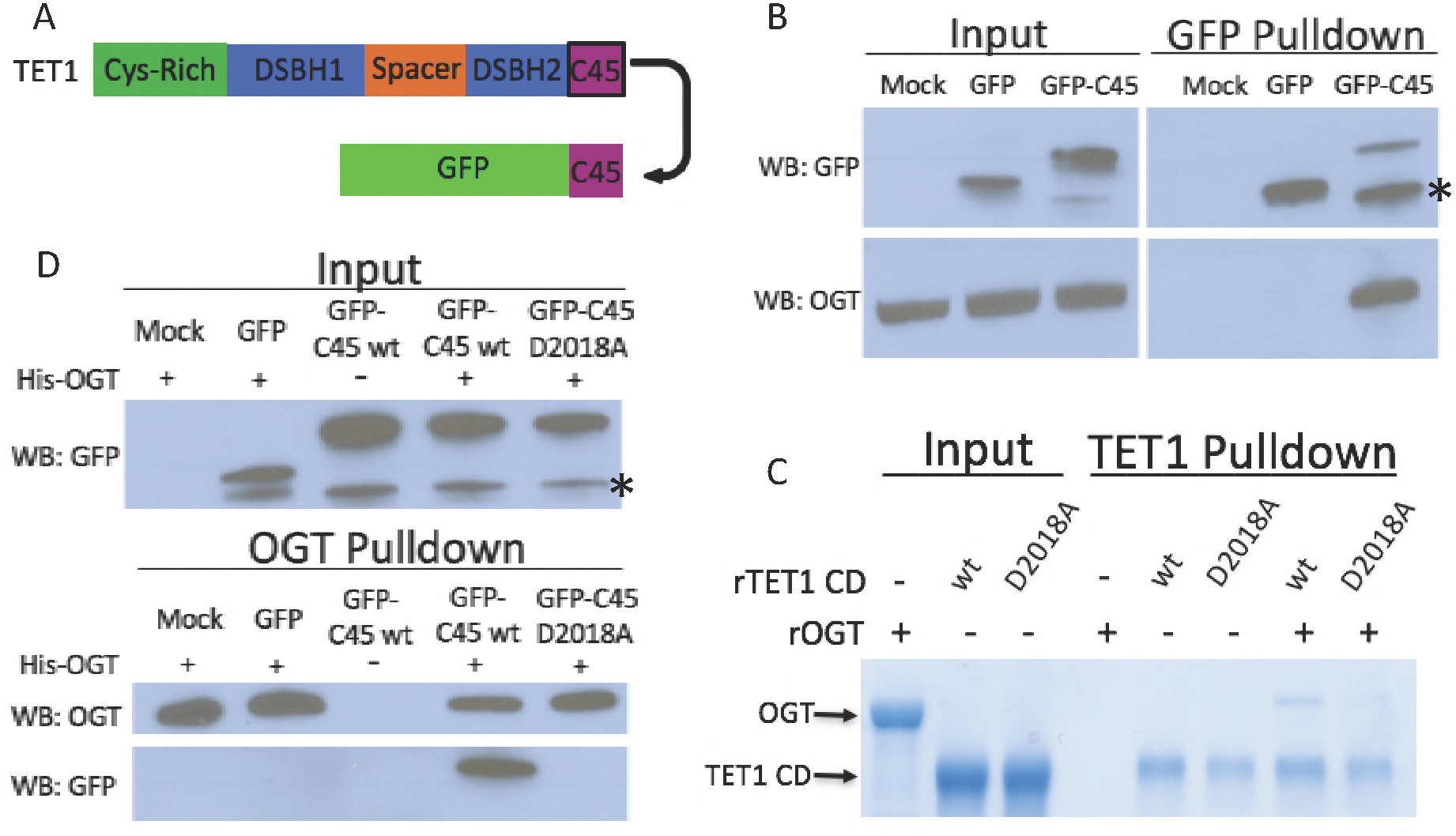
The TET1 C45 is sufficient for interaction with OGT in cells and *in vitro*. A) Schematic of the TET1 C45 fusion to the C-terminus of GFP. B) GFP and OGT western blot of inputs and GFP IPs from HEK293T cells transiently expressing GFP or GFP-TET1 C45. *Truncated GFP. C) Coomassie stained protein gel of inputs and TET1 IPs from *in vitro* binding reactions containing rOGT and rTETl CD wild type or D2018A. No UDP-GlcNAc was included in these reactions. D) GFP and OGT western blot of inputs and OGT IPs from *in vitro* binding reactions containing rOGT and *in vitro* translated GFP constructs. *Truncated GFP.

To determine if the interaction between TET1 CD and OGT is direct, we employed recombinant proteins in pulldown assays using beads conjugated to a TET1 antibody. We used recombinant human OGT (rOGT) isolated from *E. coli* and recombinant mouse TET1 catalytic domain (aa1367-2039), either wild type (rTET1 wt) or D2018A (rD2018A) purified from sf9 cells. rTET1 wt, but not beads alone, pulled down rOGT, indicating a direct interaction between these proteins (Fig. 3C). rD2018A did not pull down rOGT, consistent with our mutational analysis in cells. Then we used an *in vitro* transcription/translation extract to produce GFP and GFP-C45, incubated each with rOGT, and found that the TET1 C45 is sufficient to confer binding to rOGT (Fig. 3D). The D2018A mutation in the GFP-C45 was also sufficient to prevent rOGT binding (Fig. 3D), consistent with the behavior of TET1 CD D2018A in cells. Together these results indicate that the TET1-OGT interaction is direct and mediated by the TET1 C45.

### The D2018A mutation impairs TET1 CD stimulation by OGT

We employed the D2018A mutation to investigate the effects of perturbing the TET1-OGT interaction on rTET1 activity. rTET1 wt and rD2018A catalyzed formation of 5hmC on an *in vitro* methylated lambda DNA substrate (Fig. 4A). Incubation with rOGT and OGT’s cofactor UDP-GlcNAc resulted in *O*-GlcNAcylation of rTET1 wt but not rD2018A (Fig. 4B).

**Fig. 4:**
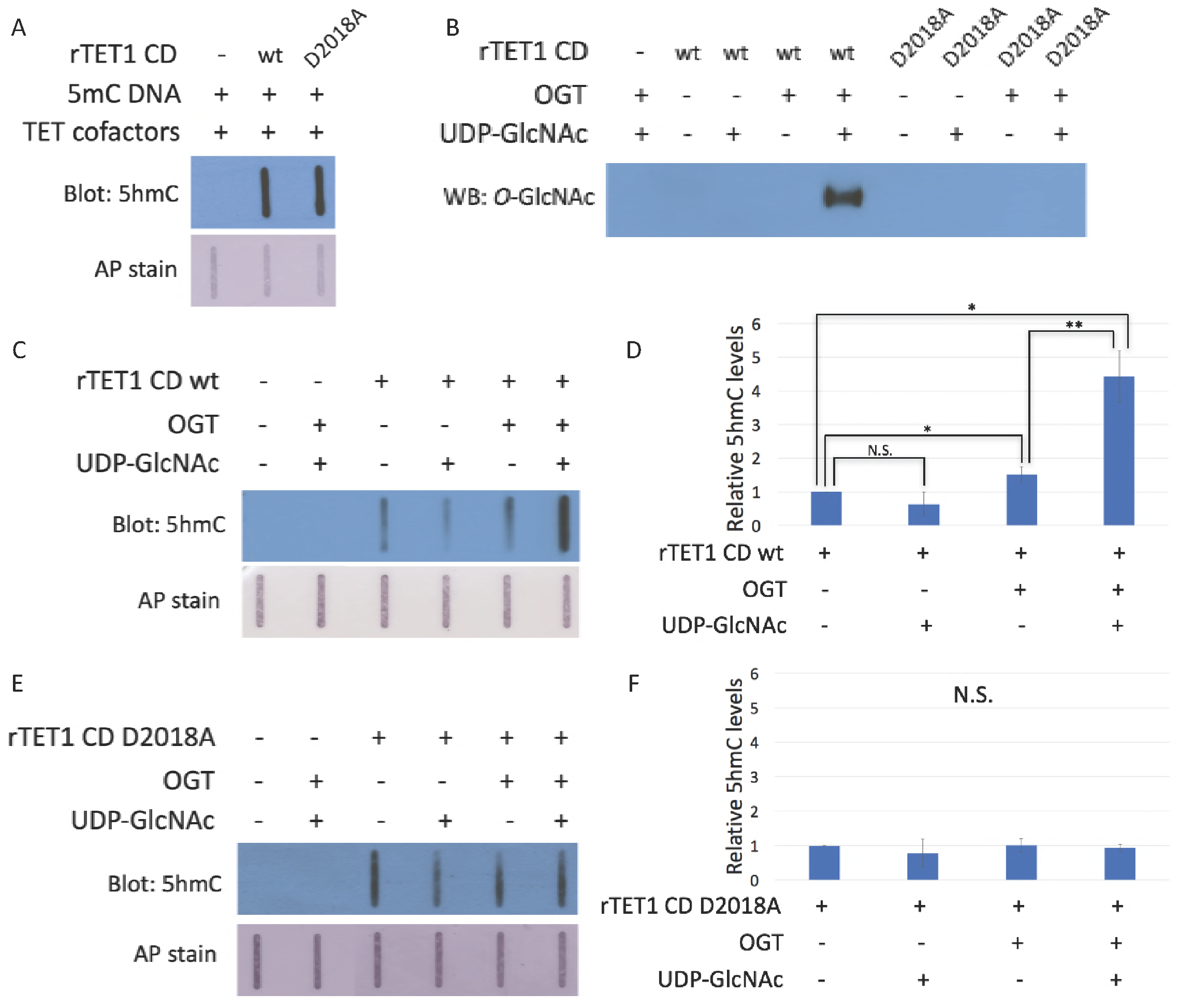
The D2018A mutation impairs TET1 CD stimulation by OGT. A) 5hmC slot blot of biotinylated 5mC containing lambda DNA from rTETl CD activity assays. Alkaline phosphatase staining was used to detect biotin as a loading control. B) Western blot for *O*-GlcNAc in *in vitro O*-GlcNAcylation reactions. C) 5hmC slot blot of biotinylated 5mC containing lambda DNA from rTETl wt activity assays. Alkaline phosphatase staining was used to detect biotin as a loading control. D) Quantification of 5hmC levels from rTETl wt activity assays. Results are from 3-5 slot blots and normalized to rTETl wt alone. E) 5hmC slot blot of biotinylated 5mC containing lambda DNA from rD2018A activity assays. Alkaline phosphatase staining was used to detect biotin as a loading control. F) Quantification of 5hmC levels from rD2018A activity assays. Results are from 3-5 slot blots and normalized to rD2018A alone. Error bars denote s.d. *P<0.01, **P<0.01, N.S. **-** not significant.

To explore whether *O*-GlcNAcylation affects TET1 CD activity, we incubated rTET1 wt and rD2018A with UDP-GlcNAc and rOGT individually or together and assessed 5hmC production. Addition of UDP-GlcNAc did not significantly affect activity of rTET1 wt or rD2018A. Incubation with rOGT alone slightly enhanced 5hmC synthesis by rTET1 wt (1.3. -1.7-fold), but not rD2018A. We observed robust stimulation of TET activity (4-5-fold) when rTET1 wt but not rD2018A was incubated with rOGT and UDP-GlcNAc (Fig. 4C-F). These results suggest that while the TET1-OGT protein-protein interaction may slightly enhance TET1’s activity, the *O*-GlcNAc modification is responsible for the majority of the observed stimulation.

### The TET-OGT interaction promotes TET1 function in the zebrafish embryo

We used zebrafish as a model system to ask whether the D2018A mutation affects TET function during development. Deletion analysis of *tet*s in zebrafish showed that Tet2 and Tet3 are the most important in development, while Tet1 contribution is relatively limited [32]. Deletion of both *tet2* and *tet3* (*tet2/3^DM^*) causes a severe decrease in 5hmC levels accompanied by larval lethality owing to abnormalities including defects in hematopoietic stem cell (HSC) production. Reduced HSC production is visualized by reductions in the transcription factor *runx1*, which marks HSCs in the dorsal aorta of wild-type embryos, but is largely absent from this region in *tet2/3^DM^* embryos. 5hmC levels and *runx1* expression are rescued by injection of human TET2 or TET3 mRNA into one-cell-stage embryos [32].

Given strong sequence conservation among vertebrate TET/Tet proteins, we asked if over expression of mouse Tet1 mRNA could also rescue HSC production in *tet2/3^DM^* zebrafish embryos and if this rescue is OGT interaction-dependent. To this end, *tet2/3^DM^* embryos were injected with wild type or D2018A mutant encoding mouse Tet1 mRNA at the one cell stage. At 30 hours post fertilization (hpf) embryos were fixed and the presence of *runx1* positive HSCs in the dorsal aorta was assessed by *in situ* hybridization (Fig. 5A). Tet1 wild type mRNA significantly increased the percentage of embryos with strong *runx1* labeling in the dorsal aorta (high *runx1*), while Tet1 D2018A mRNA failed to rescue *runx1* positive cells (Fig. 5A-B). We also performed dot blots with genomic DNA from these embryos to measure levels of 5hmC (Fig. 5C). On average, embryos injected with wild type Tet1 mRNA showed a modest but significant increase in 5hmC relative to uninjected *tet2/3^DM^* embryos, while injection of TET1 D2018A mRNA did not show a significant increase (Fig. 5D). These results suggest that the TET1-OGT interaction promotes both TET1’s catalytic activity and its ability to rescue *runx1* expression in this system.

**Fig. 5:**
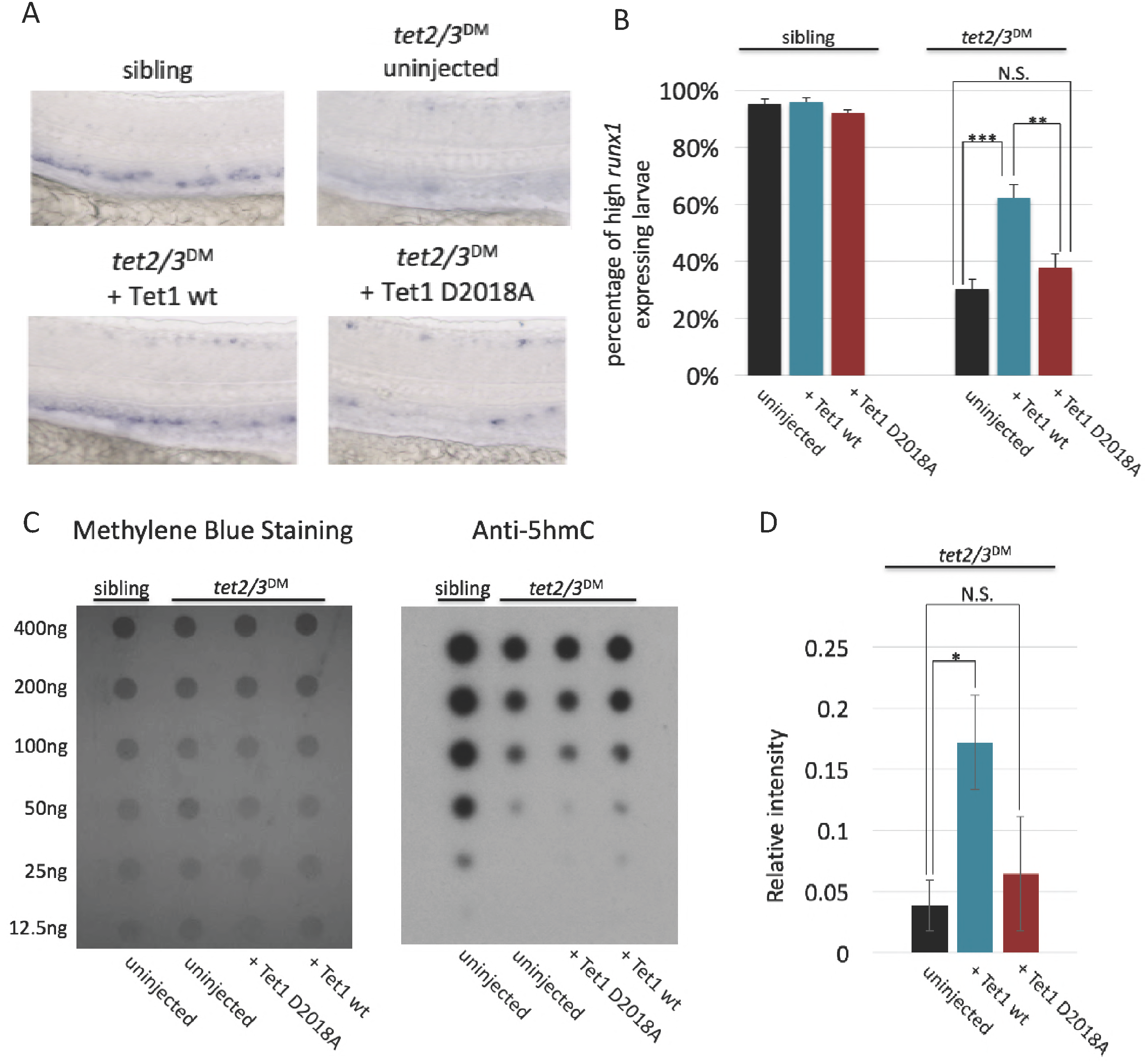
The TET1-0GT interaction promotes TET1 function in the zebrafish embryo. A) Representative images of *runxl* labeling in the dorsal aorta of wild type or *tet2/3^DM^* zebrafish embryos, uninjected or injected with mRNA encoding mouse Tetl wild type or D2018A. B) Percentage of embryos with high *runxl* expression along the dorsal aorta (*P<0.05, **P<0.01, ***P<0.001, N.S. **-** not significant). C) 5hmC dot blot of genomic DNA from wild type or *tet2/3^DM^* zebrafish embryos injected with Tetl wild type or D2018A mRNA. Methylene blue was used as a loading control. D) Quantification of 5hmC levels from 3 dot blots, normalized to methylene blue staining (*P<0.05, **P<0.01, ***P<0.001, N.S. - not significant).

### The D2018A mutation alters TET-containing complexes in mESCs

Given the defect of TET1 D2018A in the zebrafish system, we decided to explore the effect of this mutation in mammalian cells. To this end, we generated a D2018A mutation in both copies of the *Tet1* gene (Fig. 6A) in mESCs (Fig. 6 – figure supplement 1). A FLAG tag was also introduced onto the C-terminus of wild type (WT) or D2018A mutant (D2018A) TET1. We first tested whether D2018 was necessary for the TET1-OGT interaction in the context of endogenous full length TET1 in these cells. FLAG pulldowns revealed that the D2018A mutation reduced, but did not eliminate, co-IP of OGT with TET1 (Fig. 6B; Fig. 6 – figure supplement 1). Levels of 5hmC were comparable between WT and D2018A mESCs (Fig. 6C), suggesting that overall TET activity is similar.

**Fig. 6:**
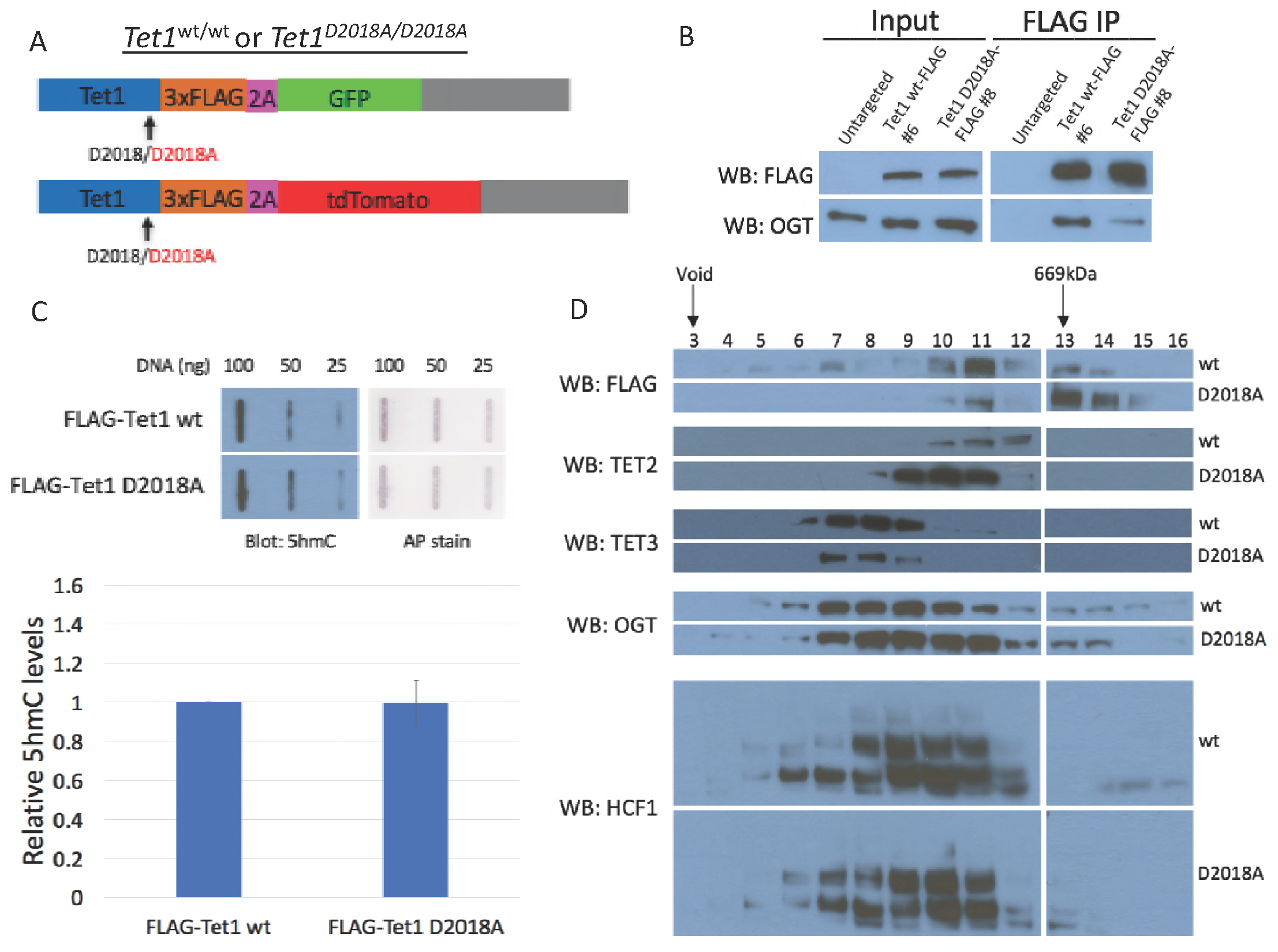
The D2018A mutation alters TET-containing complexes in mESCs. A) Schematic of WT-FLAG and D2018A-FLAG mESC lines. B) FLAG and OGT western blot of inputs and FLAG IPs from WT-FLAG and D2018A-FLAG mESCs. C) (Upper) Representative 5hmC slot blot of 25-100ng genomic DNA from WT-FLAG and D2018A-FLAG mESCs. Equal amounts of biotinylated plasmid DNA were added to each gDNA stock and diluted across the dilution series. Alkaline phosphatase staining was used to detect biotin as a loading and dilution control. (Lower) relative levels of 5hmC in WT-FLAG and D2018A-FLAG mESCs from four independent slot blots. D) Western blots for FLAG, TET2, TET3, OGT and HCF1 of nuclear extracts from WT-FLAG and D2018A-FLAG mESCs fractionated on a Superose 6 size exclusion column. Fraction numbers are indicated.

In mESCs, TETs are found in high molecular weight complexes that also contain OGT and the OGT-interacting protein HCF1 [26]. To examine whether the D2018A mutation affected these complexes, we performed size exclusion chromatography on nuclear extracts prepared from WT and D2018A mESCs (Fig. 6D). Consistent with previous reports, in WT mESCs TET1 and TET2 were found in overlapping high molecular weight (>669kDa) complexes that contain OGT and HCF1. While TET3 is the smallest of the three TETs it was found in the highest molecular weight fractions, which also contained both OGT and HCF1. In D2018A mESCs all three TET-containing complexes were altered. Although the total amount of FLAG-TET1 was comparable between WT and D2018A cells (Fig. 6B), in D2018A mESCs the amount of FLAG-TET1 in high molecular weight fractions was reduced (Fig. 6D). This reduction coincided with an increase in abundance of FLAG-TET1 in lower molecular weight fractions that contained much less OGT and HCF1. In contrast, TET2 increased in abundance and shifted to higher molecular weight fractions, while TET3 decreased in abundance but remained in the same fractions (Fig. 6D). These results suggest that the normal interaction between TET1 and OGT is necessary for the proper distribution of TET1, TET2, and TET3 in high molecular weight complexes. The increase in the amount of TET2 in D2018A mESCs may explain why bulk 5hmC levels were not appreciably affected by this mutation (Fig 6C).

### The D2018A mutation alters 5mC distribution and gene expression

To determine whether these alterations in TET-containing high molecular weight complexes affected gene expression, we compared WT and D2018A mESCs using RNA-seq. Of the roughly 8800 expressed genes (FDR <0.1), we found over 2000 genes whose expression changed by 2-fold or more in D2018A cells compared to WT (596 upregulated genes and 1639 downregulated genes) (Fig. 7A, Supplementary File 4). These results show that a single amino acid substitution in TET1 causes a substantial change in global gene expression.

**Fig. 7:**
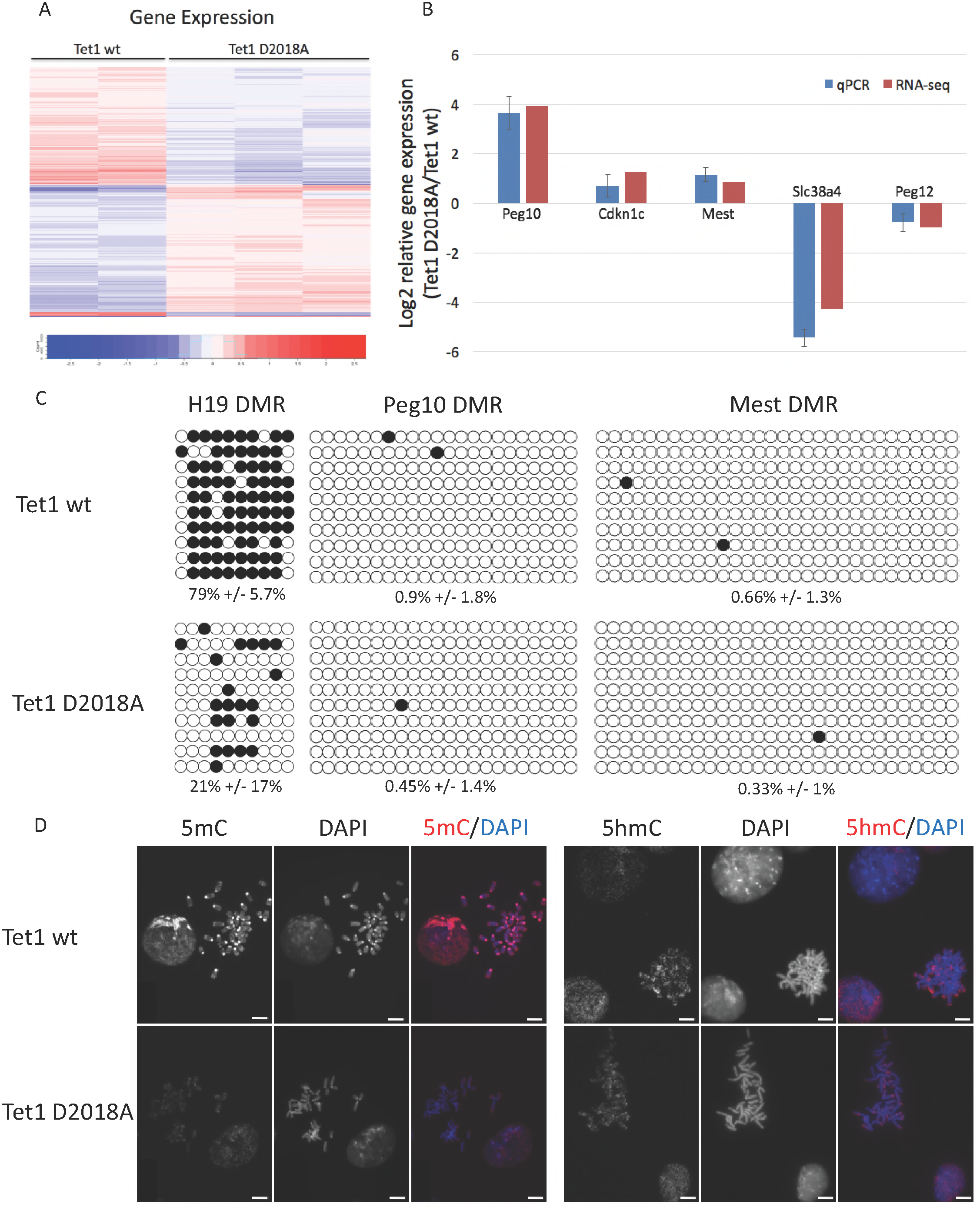
The D2018A mutation alters 5mC distribution and gene expression. A) Heatmap depicting gene expression changes between Tetl wt and D2018A mESCs. B) RT-qPCR analysis of 5 selected imprinted genes from the RNA-seq dataset. Each reaction was performed in triplicate. Error bars represent s.d. C) Targeted bisulfite analysis of DMRs associated with 3 imprinted genes. Filled circles depict 5mC or BhmC, empty circles depict unmodified C. Error represents s.d. D) Immunofluorescence staining for 5mC and 5hmC on chromosome spreads. Scale bar: lOum.

In mouse development TET1 is necessary for normal expression of many imprinted genes[36], prompting us to examine this class of genes. Of the 35 imprinted genes with detectable expression (FDR <0.1) in either WT or D2018A mESCs, 14 changed expression by 2fold or more and an additional 8 changed expression by 1.5-2-fold (Supplementary File 5). RT-qPCR for selected imprinted genes in WT and D2018A mESCs confirmed the gene expression changes found by RNA-seq (Fig. 7B). These data suggest that imprinted genes may be subject to regulation by TET1 and OGT in mESCs.

Since TETs act to remove DNA methylation, we wondered whether the changes in imprinted gene expression in D2018A mESCs might be due to changes in the methylation status of differentially methylated regions (DMRs). We therefore performed targeted bisulfite analysis of DMRs associated with three imprinted genes, *H19*, *Peg10*, and *Mest*, which were upregulated in D2018A mESCs compared to WT. We found that the *H19* DMR was heavily methylated in WT cells (79% +/- 5.7%) but significantly less methylated in D2018A cells (21% +/-17%) (Fig. 7C), consistent with the very large (∼30-fold) increase in expression of *H19* in D2018A cells compared to WT. In contrast, the *Peg10* and *Mest* DMRs were almost completely unmethylated in both cell types (Fig. 7C), indicating that large changes in DMR methylation do not account for the altered expression of these imprinted genes.

To examine whether regions other than DMRs exhibited altered cytosine modifications, we performed immunofluorescence staining for 5mC and 5hmC on chromosome spread preparations from WT and D2018A mESCs (Fig. 7D). Although 5hmC staining was comparable between the two cell lines, we observed a striking difference in the distribution of 5mC. While WT cells showed enrichment of 5mC staining at pericentric heterochromatin, no such enrichment was observed in D2018A cells. These analyses of cytosine modifications at imprinted gene DMRs and pericentric heterochromatin indicate that the D2018A mutation has a substantial impact on 5mC abundance and distribution, as well as gene expression.

## Discussion

### A unique OGT interaction domain?

We identified a 45-amino acid domain of TET1 that is both necessary and sufficient for binding of OGT. To our knowledge, this is the first time that a small protein domain has been identified that confers stable binding to OGT. The vast majority of OGT targets do not bind to OGT tightly enough to be detected in co-IP experiments, suggesting that OGT’s interaction with TET proteins is unusually strong. For determination of the crystal structure of the human TET2 catalytic domain in complex with DNA, the corresponding C-terminal region was deleted [37], suggesting that it may be unstructured. When bound to OGT this domain may become structured, and structural studies of OGT bound to C45 could shed light on what features make this domain uniquely able to interact stably with OGT and how OGT may stimulate TET1 activity.

An alternative or additional role for the stable TET-OGT interaction may be recruitment of OGT to chromatin by TET proteins. Loss of TET1 causes loss of OGT from chromatin [26] and induces similar changes in transcription in both wild-type mESCs and mESCs lacking DNA methylation [38]. This raises the possibility that TET proteins may recruit OGT to chromatin to regulate gene expression independent of 5mC oxidation. Consistent with this possibility, OGT modifies many transcription factors and chromatin regulators in mESCs [39](Fig. 8). Thus it may be that the stable TET1-OGT interaction promotes both regulation of TET1 activity by *O*-GlcNAcylation as well as recruitment of OGT to chromatin. Notably, our results show that TET1 D2018A does not rescue 5hmC levels in *tet2/3^DM^* zebrafish embryos to the same extent as the wild type protein, suggesting that at least part of the role of the TET1-OGT interaction *in vivo* is regulation of TET1 activity.

**Fig. 8:**
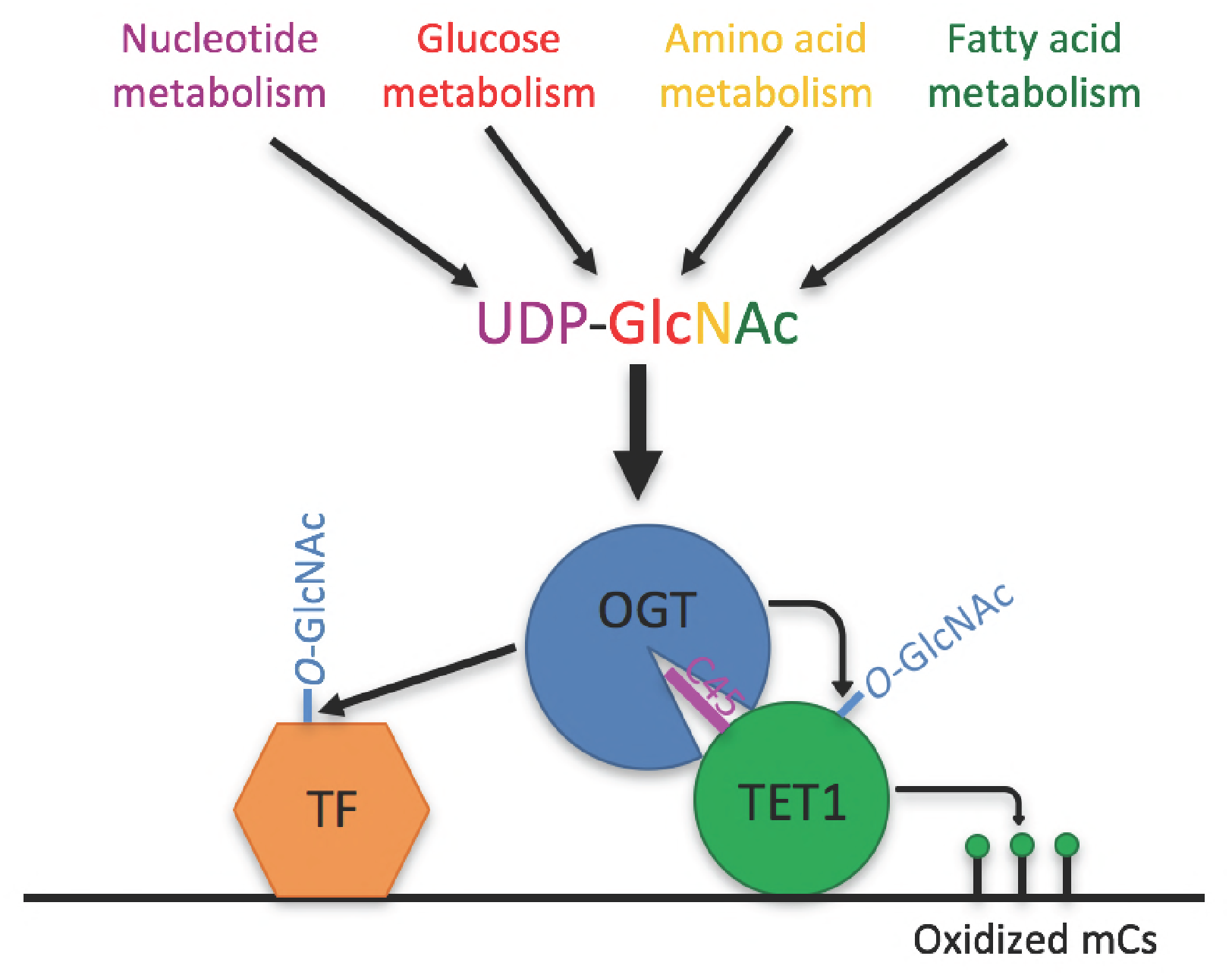
Model. Model showing two roles of the TET1;OGT interaction in regulation of gene expression. OGT’s activity is regulated by the abundance of its cofactor UDP;GlcNAc, whose synthesis has inputs from nucleotide, glucose, amino acid, and fatty acid metabolism. OGT (blue circle) binds to TET1 (large green circle) via the TET1 C45 (purple line). OGT modifies TET1 and regulates its catalytic activity (small green circles representing modified cytosines). At the same time, TET1 binding to DNA brings OGT into proximity of other DNA;bound transcription factors (orange hexagon), which OGT also modifies and regulates.

### OGT stimulation of TET activity

Our results show for the first time that OGT can modify a TET protein *in vitro*, and that *O*-GlcNAcylation stimulates the activity of a TET protein *in vitro*. We have identified 8 sites of *O*-GlcNAcylation within the TET1 CD (data not shown), which precludes a simple analysis of which sites are important for stimulation. It is unclear how many sites are important for TET1 function, as it is possible that the unusually stable interaction between OGT and TET1 allows OGT to nonspecifically modify serine/threonine resides on TET1. Detailed studies of individual sites of modification will be required to resolve this question.

Our data are also consistent with a role for OGT in TET1 regulation in cells and *in vivo*. OGT also directly interacts with TET2 and TET3, suggesting that it may regulate all three TET proteins. Notably, although all three TETs catalyze the same reaction, they show a number of differences that are likely to determine their biological role. Different TET proteins are expressed in different cell types and at different stages of development [40-43]. TET1 and TET2 appear to target different genomic regions [44] and to promote different pluripotent states in mESCs [45]. The mechanisms responsible for these differences are not well understood. We suggest that OGT is a strong candidate for regulation of TET enzymes.

### Regulation of TETs by OGT in development

Our result that wild type TET1 mRNA, but not TET1 mRNA carrying a mutation that can impair interaction with OGT, can rescue *tet2/3^DM^* zebrafish suggests that OGT regulation of TET enzymes may play a role in development. The importance of both TET proteins and OGT in development has been thoroughly established. Zebrafish lacking *tet2* and *tet3* die as larvae [32], and knockout of *Tet* genes in mice yields developmental phenotypes of varying severities, with knockout of all three *Tet*s together being embryonic lethal [41,42,46,47]. Similarly, OGT is absolutely essential for development in mice [48] and zebrafish [49], though its vast number of targets have made it difficult to narrow down more specifically why OGT is necessary. Our results suggest that TETs are important OGT targets in development.

### The TET1-OGT interaction regulates TET-containing complexes and gene expression in mESCs

The D2018A mutation reduced the TET1-OGT interaction in mESCs and altered all 3 TET containing high molecular weight complexes. While these changes did not correlate with alterations in bulk 5hmC levels, the distribution of 5mC was altered. The region of TET1 that is necessary and sufficient for interaction with OGT is highly conserved with the other TETs and perturbing the interaction between OGT and TET1 altered the abundance of TET2 and TET3 in high molecular weight complexes. Together these data suggest that OGT may be equilibrating between the three TET-containing complexes. The size of the complexes in which TETs are found (>670kDa) are larger than would be expected if the only components are a TET protein, OGT, and HCF1, suggesting that additional proteins or more than one molecule of OGT, HCF1, or TET are present. A thorough study of the factors that comprise these complexes, as well as how the TET1 D2018A mutation alters the architecture of these complexes and the epigenetic status of the genome will yield valuable insights into how the TET1-OGT interaction regulates gene expression in mESCs.

The D2018A mutation caused a large increase in the levels of TET2, which may explain why bulk 5hmC levels are unaltered when the TET1-OGT interaction is decreased. TET1 and TET2 regulate different genomic regions in mESCs[44], and redistribution of TET2 to TET1 targets may contribute to the altered distribution of 5mC and gene expression seen in the D2018A mESCs. The magnitude of gene expression changes (nearly one quarter of genes changed 2-fold or more) and striking alteration in 5mC distribution induced by a single amino acid substitution demonstrates the importance of the TET1-OGT interaction in regulation of the transcriptome and epigenome. Further study of how 5mC/5hmC levels and distribution are controlled by the TET1-OGT interaction will provide insight into how this nutrient-sensing post-translational modification enzyme can regulate the epigenome.

### A connection between metabolism and the epigenome

OGT has been proposed to act as a metabolic sensor because its cofactor, UDP-GlcNAc, is synthesized via the hexosamine biosynthetic pathway (HBP), which is fed by pathways metabolizing glucose, amino acids, fatty acids, and nucleotides [24]. UDP-GlcNAc levels change in response to flux through these pathways [50-52], leading to the hypothesis that OGT activity may vary in response to the nutrient status of the cell. Thus the enhancement of TET1 activity by OGT and the significant overlap of the two enzymes on chromatin [26] suggest a model in which OGT may regulate the epigenome in response to nutrient status by controlling TET1 activity (Fig. 8).

## Acknowledgments

We thank Miguel Ramalho-Santos for the FLAG-TET1 CD plasmid and Suzanne Walker for the His-OGT plasmid. We thank Leeanne Goodrich, Richard Yan, Myles Hochman, Sy Redding, Walter Eckalbar, and Andrea Barczak for technical assistance. We thank all members of the Panning lab for valuable ideas and discussion. This work was supported by R01 GM088506 (BP), NCI grant P30 CA008748 (MG), and funding from the Geoffrey Beene Cancer Research Center of Memorial Sloan-Kettering Cancer Center (MG). JH was supported by the California Institute for Regenerative Medicine Predoctoral Fellowship TG2-01153.

---

**Fig. 1 supplement 1:**
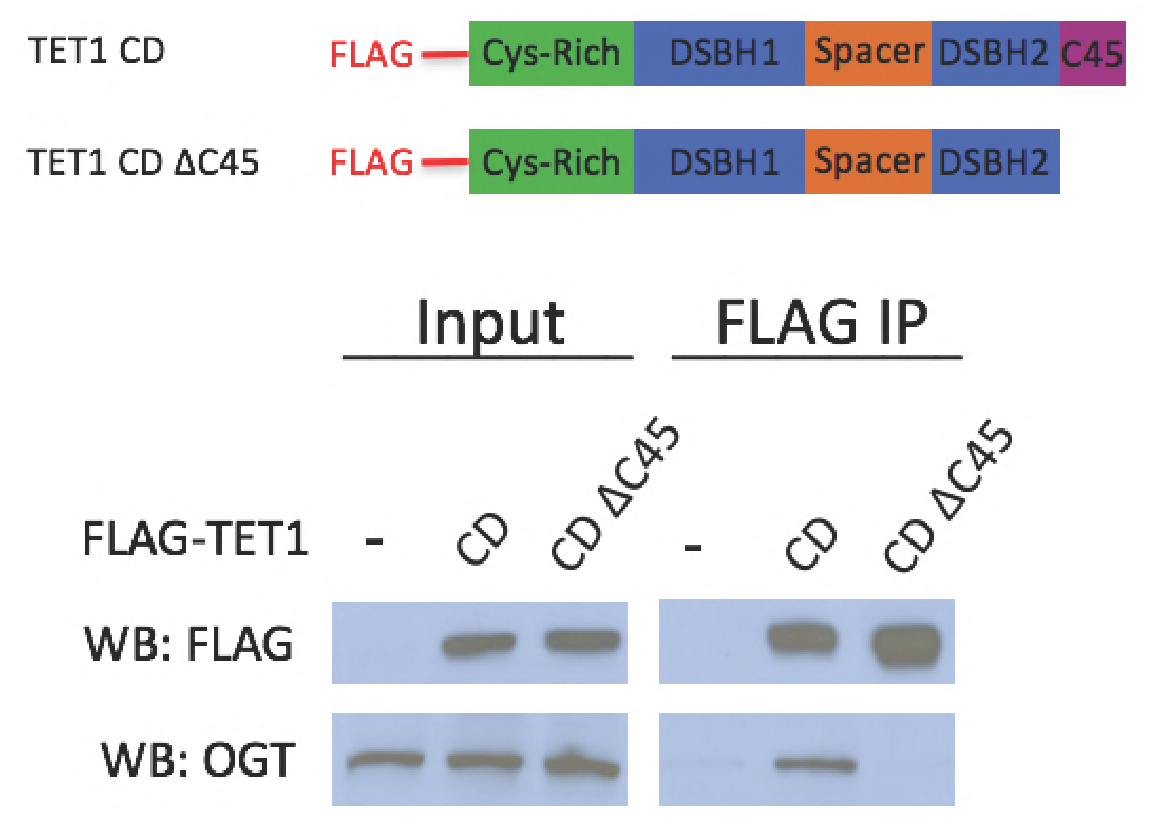
TET1 C45 is necessary for interaction with endogenous OGT. FLAG and OGT western blot of inputs and FLAG IPs from HEK293T cells transiently expressing FLAG-TET1 CD or FLAG-TET1 CD AC45 (diagrammed in the upper panel).

**Fig. 6 supplement 1:**
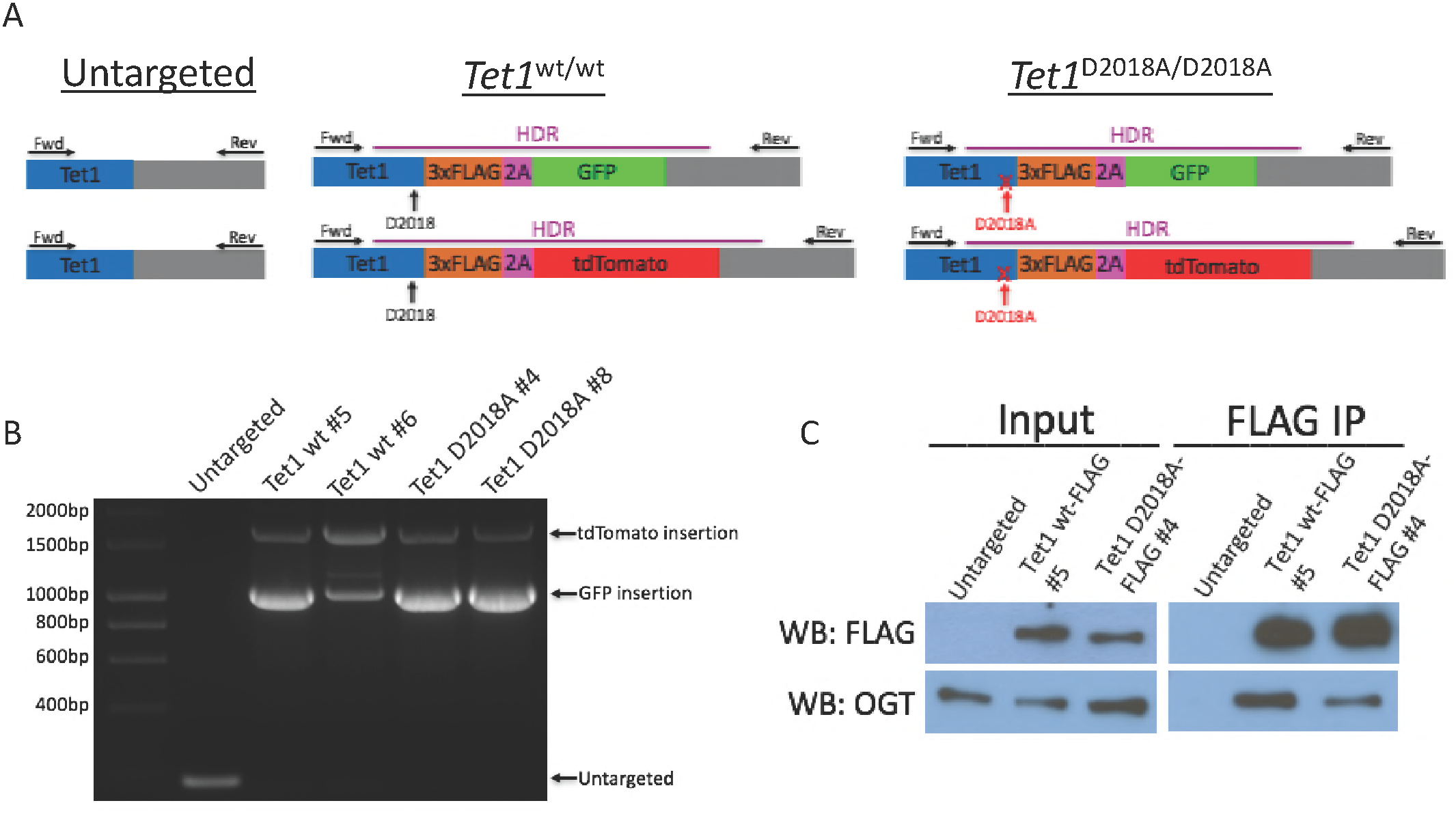
Generation of mESC lines. A) Schematic of mESC lines. DNA encoding a 3xFLAG tag was added to the 3’ end of both alleles of *Tetl,* followed by a 2A sequence and a fluorescent protein (GFP or tdTomato). The 2A sequence causes ribosome skipping, resulting in separate translation of TETl-3xFLAG and 2A-GFP or 2A-tdTomato. Purple line: template used for homology-directed repair (HDR). Horizontal arrows: primers used for PCR genotyping. Vertical arrows: D2018 residue. B) PCR genotyping of independently derived, clonal, targeted mESC lines using primers indicated in A. C) FLAG and OGT western blot of inputs and FLAG IPs from another pair of WT-FLAG and D2018A-FLAG mESCs.

**Supplementary File 1A:**
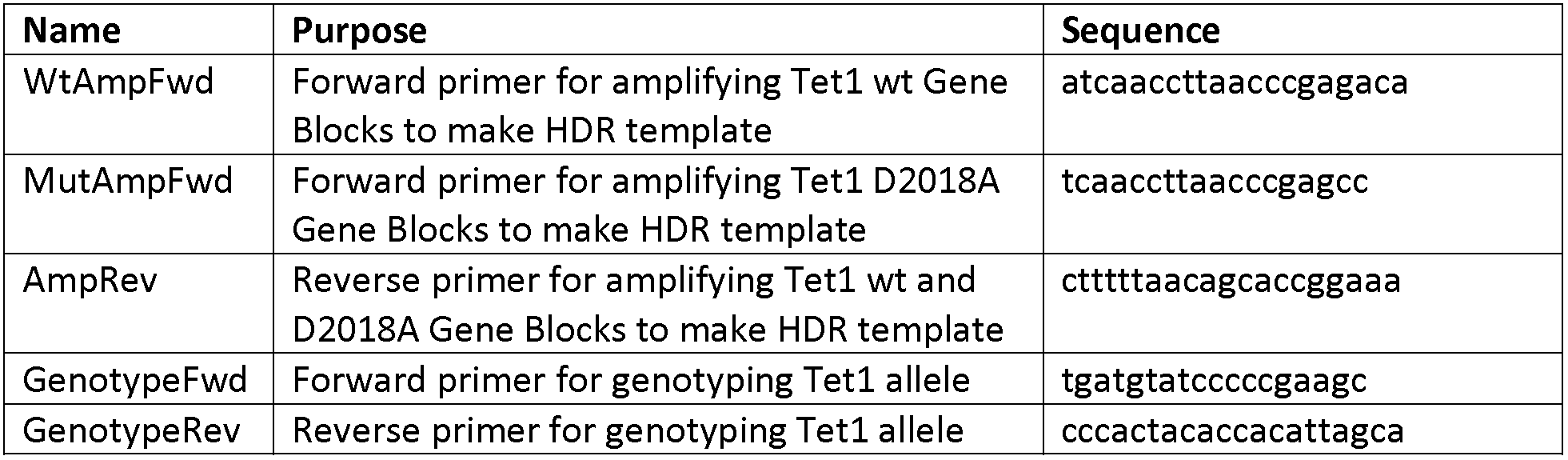
**Primers used for creating and genotyping mESC lines**

**Supplementary File 1B:**
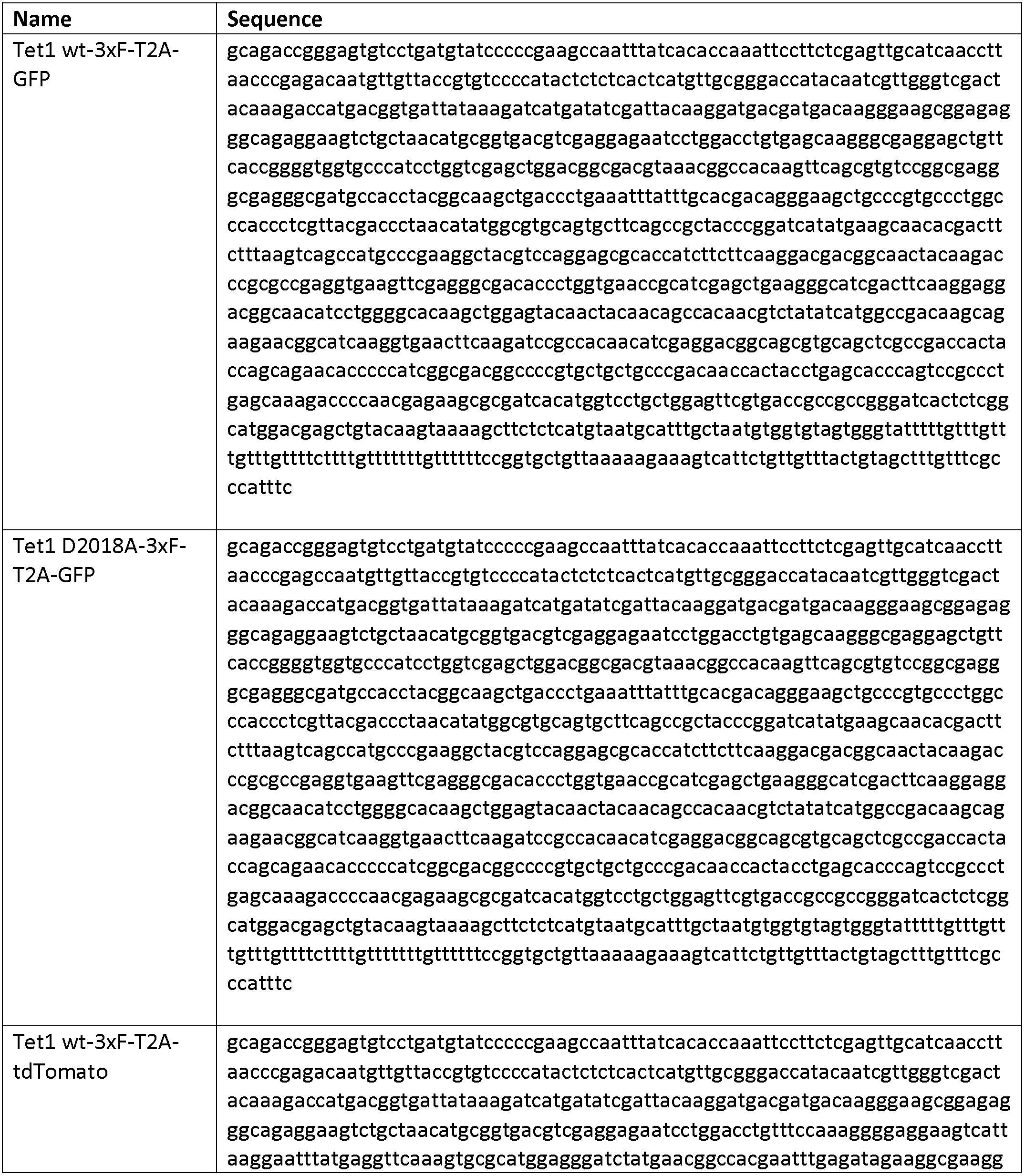
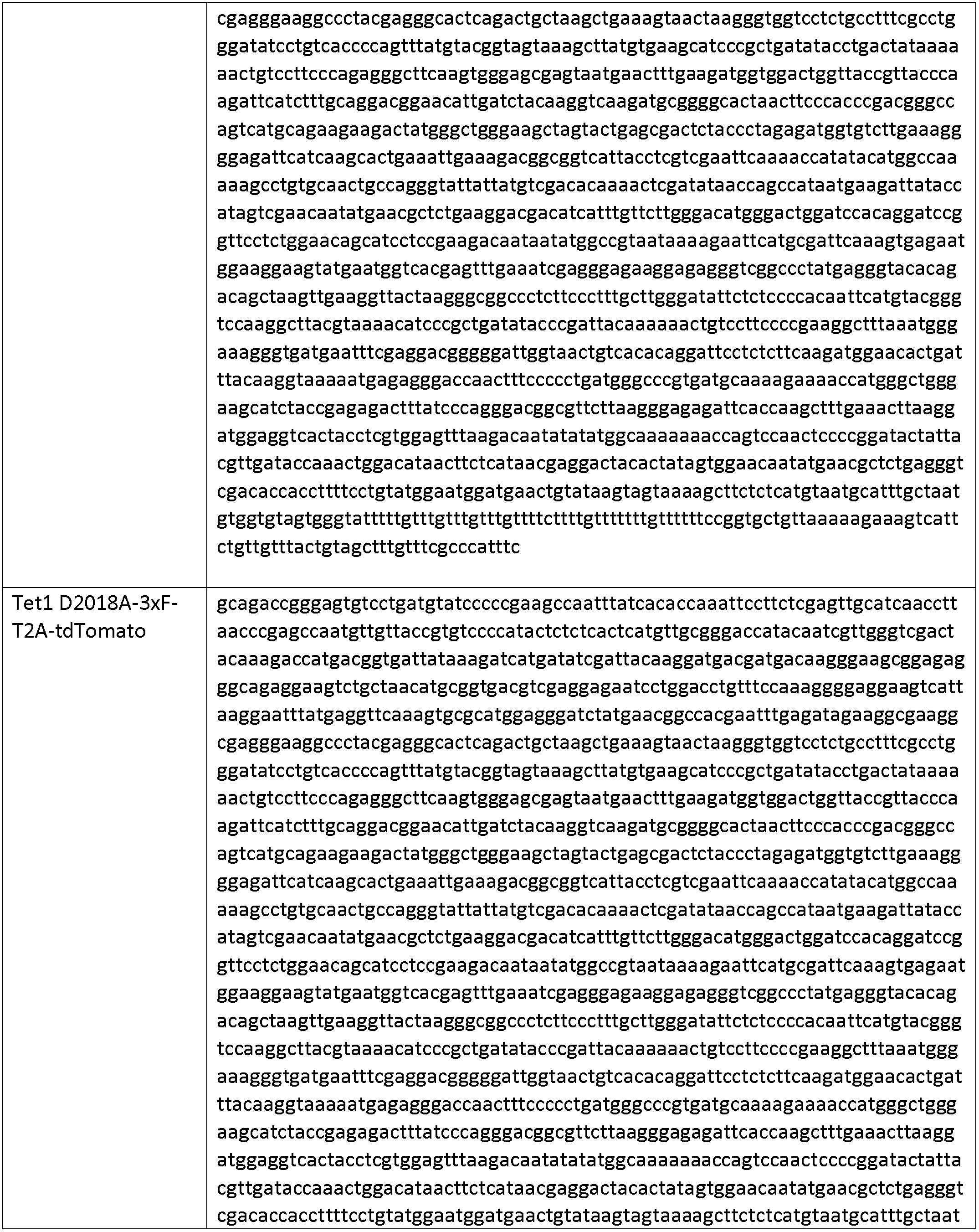
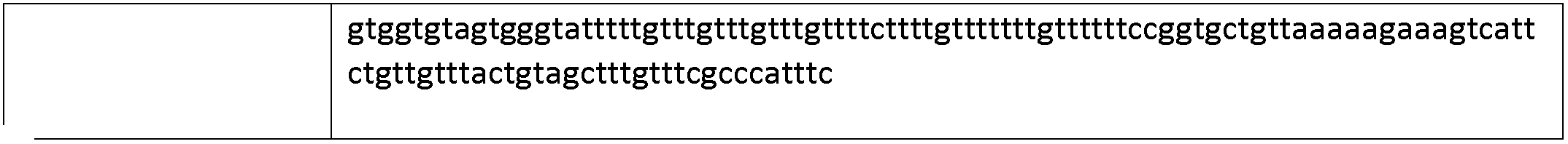
**Gene blocks amplified to make HDR templates**

**Supplementary File 2A:**
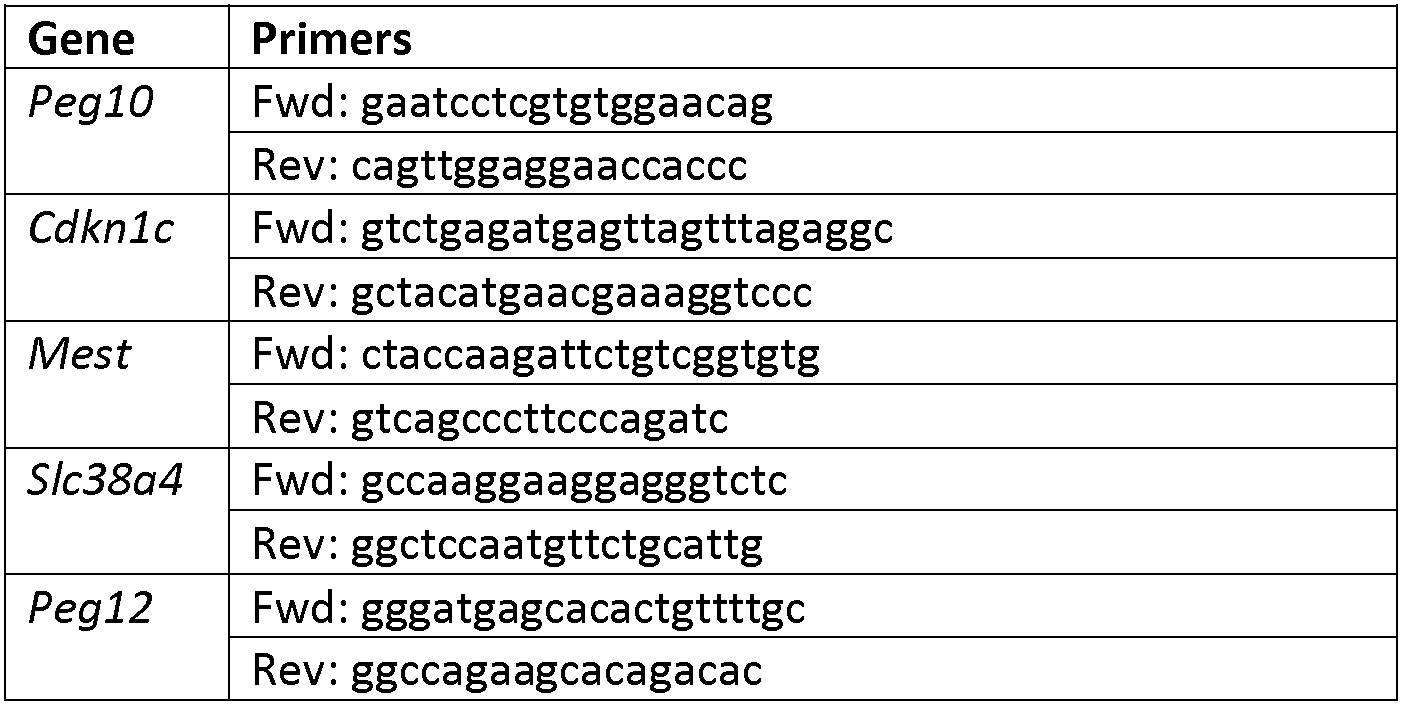
**Primers used for qPCR**

**Supplementary File 2B:**
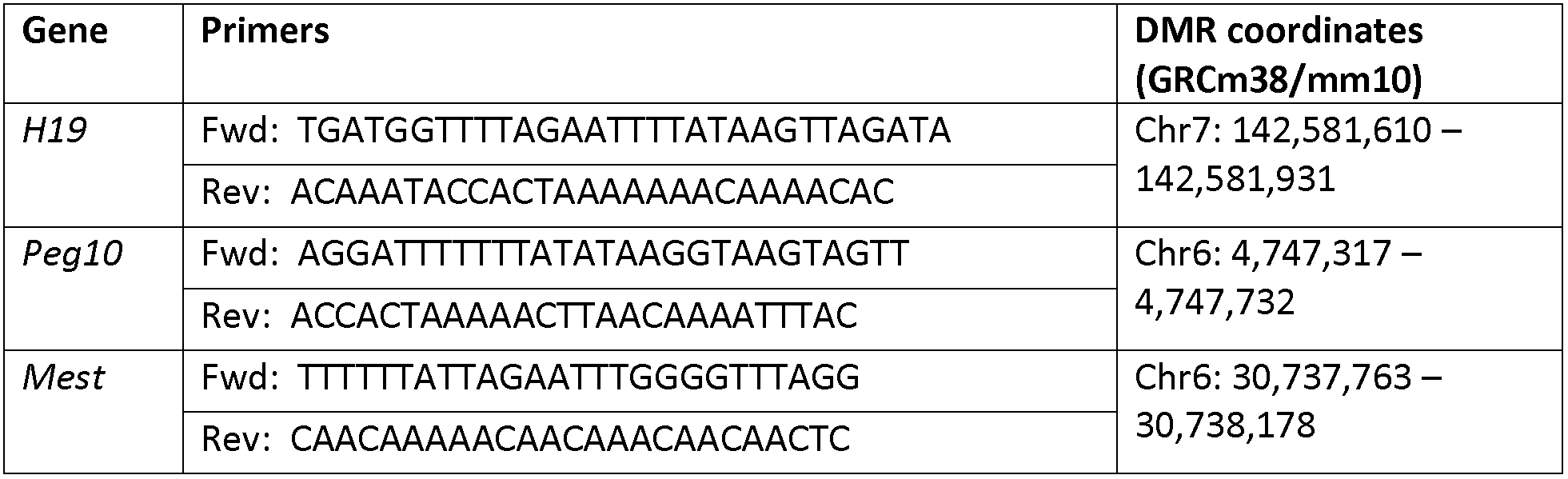
**Primers and DMRs used in bisulfite analysis** Phage lambda control primers: Fwd: AGTTTGTTATTGTTAGGAAAGTGGTAAA Rev: TCAACCTAAATCATTAAAACCTACC

**Supplementary File 3:**
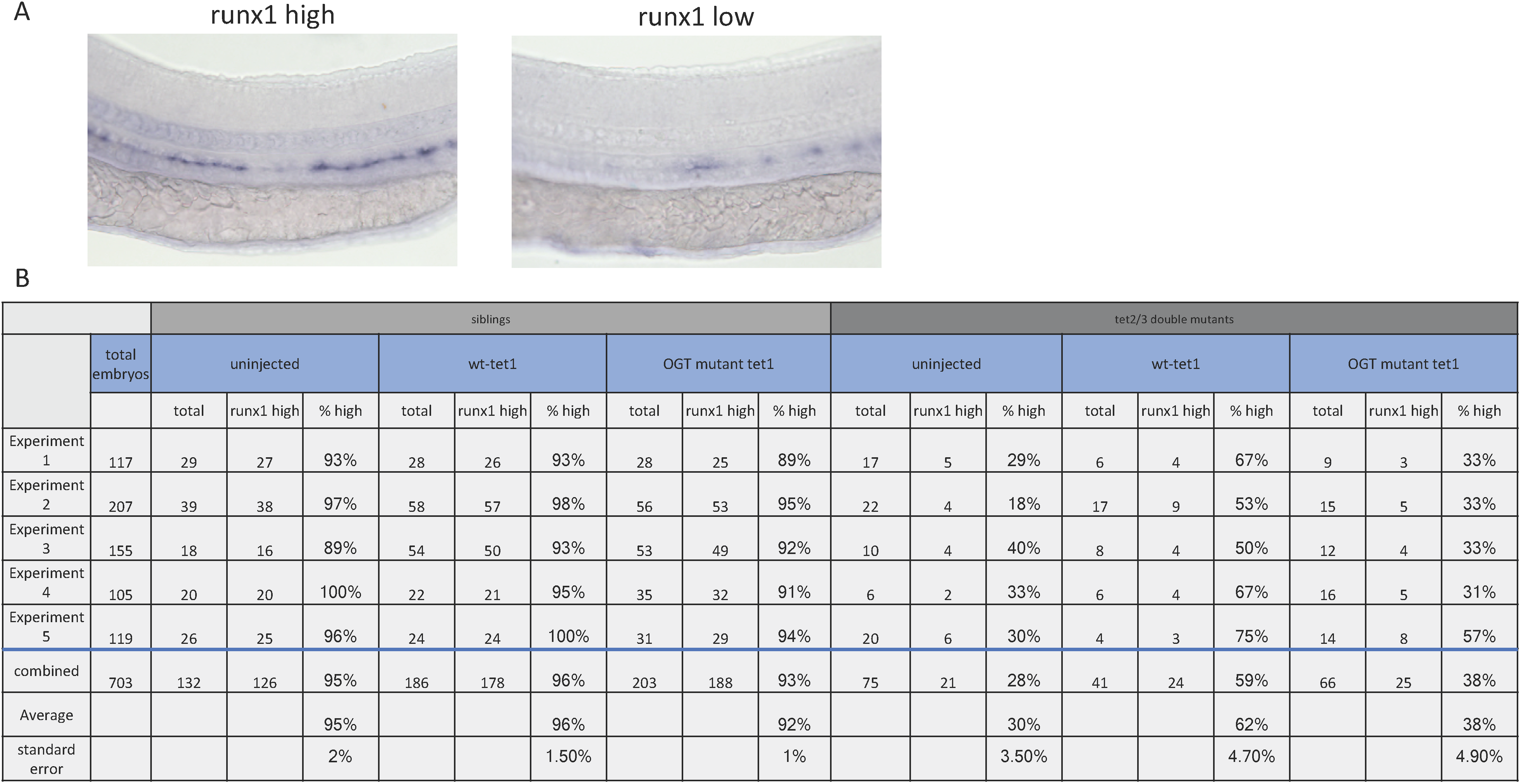
**Analysis of zebrafish larvae** A) Representative images of larvae with high and low *runxl* expression. B) Embryo numbers and scoring for all 5 biological replicates.

**Supplementary File 5:**
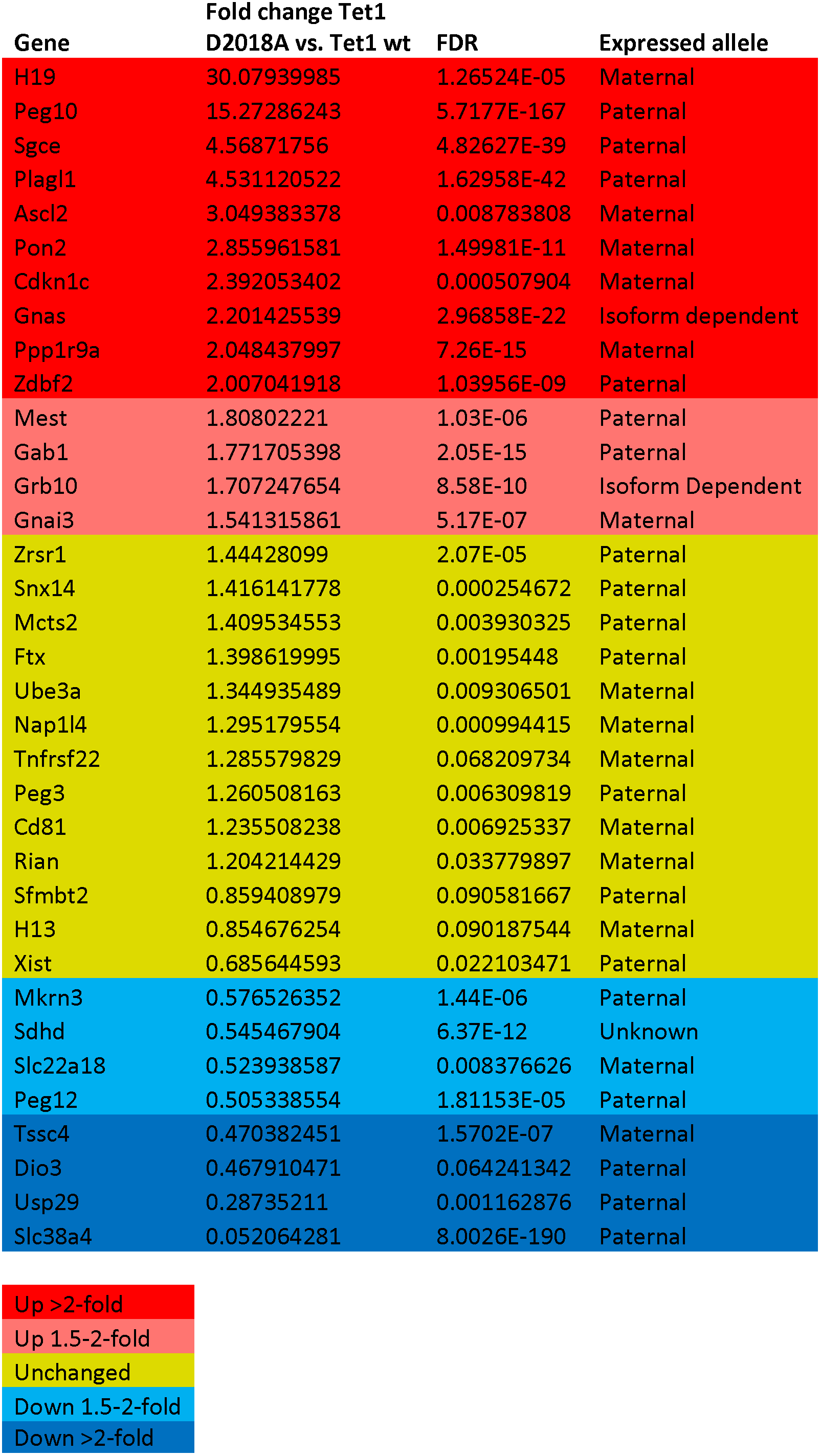
**Imprinted genes expressed in WT-FLAG and D2018A-FLAG mESCs by RNA-seq (FDR<0.1)**

| Gene | Fold change Tet1 |  | Expressed allele |
| --- | --- | --- | --- |
|  | D2018A vs. Tet1 wt | FDR |  |
| H19 | 30.07939985 | 1.26524E-05 | Maternal |
| Peg10 | 15.27286243 | 5.7177E-167 | Paternal |
| Sgce | 4.56871756 | 4.82627E-39 | Paternal |
| Plagl1 | 4.531120522 | 1.62958E-42 | Paternal |
| Ascl2 | 3.049383378 | 0.008783808 | Maternal |
| Pon2 | 2.855961581 | 1.49981E-11 | Maternal |
| Cdkn1c | 2.392053402 | 0.000507904 | Maternal |
| Gnas | 2.201425539 | 2.96858E-22 | Isoform dependent |
| Ppp1r9a | 2.048437997 | 7.26E-15 | Maternal |
| Zdbf2 | 2.007041918 | 1.03956E-09 | Paternal |
| Mest | 1.80802221 | 1.03E-06 | Paternal |
| Gab1 | 1.771705398 | 2.05E-15 | Paternal |
| Grb10 | 1.707247654 | 8.58E-10 | Isoform Dependent |
| Gnai3 | 1.541315861 | 5.17E-07 | Maternal |
| Zrsr1 | 1.44428099 | 2.07E-05 | Paternal |
| Snx14 | 1.416141778 | 0.000254672 | Paternal |
| Mcts2 | 1.409534553 | 0.003930325 | Paternal |
| Ftx | 1.398619995 | 0.00195448 | Paternal |
| Ube3a | 1.344935489 | 0.009306501 | Maternal |
| Nap1l4 | 1.295179554 | 0.000994415 | Maternal |
| Tnfrsf22 | 1.285579829 | 0.068209734 | Maternal |
| Peg3 | 1.260508163 | 0.006309819 | Paternal |
| Cd81 | 1.235508238 | 0.006925337 | Maternal |
| Rian | 1.204214429 | 0.033779897 | Maternal |
| Sfmbt2 | 0.859408979 | 0.090581667 | Paternal |
| H13 | 0.854676254 | 0.090187544 | Maternal |
| Xist | 0.685644593 | 0.022103471 | Paternal |
| Mkrn3 | 0.576526352 | 1.44E-06 | Paternal |
| Sdhd | 0.545467904 | 6.37E-12 | Unknown |
| Slc22a18 | 0.523938587 | 0.008376626 | Maternal |
| Peg12 | 0.505338554 | 1.81153E-05 | Paternal |
| Tssc4 | 0.470382451 | 1.5702E-07 | Maternal |
| Dio3 | 0.467910471 | 0.064241342 | Paternal |
| Usp29 | 0.28735211 | 0.001162876 | Paternal |
| Slc38a4 | 0.052064281 | 8.0026E-190 | Paternal |

|  |
| --- |
| Up >2-fold |
| Up 1.5-2-fold |
| Unchanged |
| Down 1.5-2-fold |
| Down >2-fold |

